# Single-molecule spectroscopy reveals dynamic allostery mediated by the substrate-binding domain of a AAA+ machine

**DOI:** 10.1101/2020.09.13.295345

**Authors:** Marija Iljina, Hisham Mazal, Pierre Goloubinoff, Inbal Riven, Gilad Haran

## Abstract

ClpB is an ATP-dependent protein disaggregation machine that is activated on demand by co-chaperones and by aggregates caused by heat shock or mutations. The regulation of ClpB’s function is critical, since its persistent activation is toxic *in vivo*. Each ClpB molecule is composed of an auxiliary N-terminal domain (NTD), an essential regulatory middle domain (MD) that activates the machine by tilting, and two nucleotide-binding domains that are responsible for ATP-fuelled substrate threading. The NTD is generally thought to serve as a substrate-binding domain, which is commonly considered to be dispensable for ClpB’s activity, and is not well-characterized structurally due to its high mobility. Here we use single-molecule FRET spectroscopy to directly monitor the real-time dynamics of ClpB’s NTD and reveal its involvement in novel allosteric interactions. We find that the NTD fluctuates on a microsecond timescale and, unexpectedly, shows little change in conformational dynamics upon binding of a substrate protein. During its fast motion, the NTD makes crucial contacts with the regulatory MD, directly affecting its conformational state and thereby influencing the overall ATPase and unfolding activity of this machine. Moreover, we also show that the NTD mediates signal transduction to the nucleotide-binding domains through conserved residues. The two regulatory pathways revealed here enable the NTD to suppress the MD in the absence of protein substrate, and to limit ATPase and disaggregation activities of ClpB. The use of multiple parallel allosteric pathways involving ultrafast domain motions might be common to AAA+ molecular machines to ensure their fast and reversible activation.

---

sAAA+ proteins are large molecular machines that convert the energy of ATP hydrolysis into mechanical work (1). Their diverse biological functions are tightly regulated through multiple allosteric interactions between their domains and subunits (2). The bacterial heat-shock protein ClpB is a AAA+ molecular machine that works in collaboration with the DnaK chaperone system (DnaK, DnaJ and GrpE) to rescue proteins from aggregates and is essential for conferring thermotolerance (3, 4). In its functional form, ClpB is a ring-shaped homohexamer that disaggregates substrate proteins by actively threading them through its central channel in an ATP-dependent process (5). Each monomer of ClpB comprises several domains (6) (Fig. 1a): a flexibly-connected N-terminal domain (NTD), two ATP-binding domains (NBD1 and NBD2), and a regulatory middle domain (MD) that binds to the co-chaperone DnaK. The NTD is a globular alpha-helical domain, with a structure that is highly conserved across the ClpA, ClpB and ClpC subfamilies (7) and in eukaryotic homologues (8). In *Escherichia coli (E. coli)*, a truncated variant lacking the NTD, ΔNClpB, is naturally co-expressed as a minor product together with the full-length ClpB (9, 10) and was reported to contribute to bacterial thermotolerance (11). The biological relevance of the truncated variant, compared to the full-length protein, is not clear.

**Figure 1.**
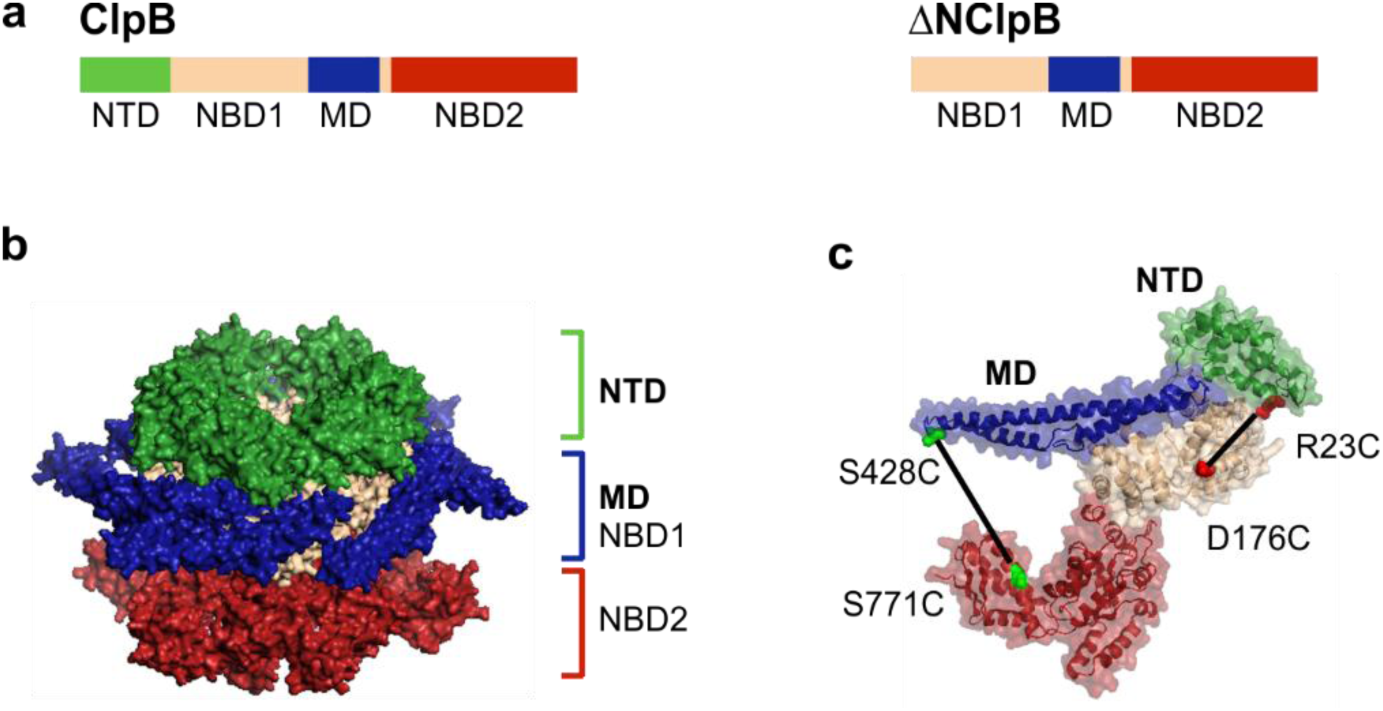
Structure of ClpB. **(a)** The sequence of domains in the full-length ClpB. A truncated, naturally occurring variant of ClpB, ΔNClpB, lacks the NTD. **(b)** Structure of the ClpB hexamer with the NTDs highlighted in green (PDB: 1QVR) (6, 37). **(c)** Structure of the ClpB monomer with the NTD shown in green and the MD in blue. Pairs of residues mutated to cysteines for the incorporation of fluorescent dyes are highlighted: R23C and D176C (in red) for the characterization of NTD dynamics; S428C and S771 (in green) for the measurement of MD dynamics.

ClpB has a complex multi-step activation mechanism (12, 13). It was shown that for its full activation, characterized by high ATPase activity and enhanced disaggregation and refolding rates, both substrate binding and the presence of the DnaK system are required (12). The MDs of ClpB form a belt surrounding the top NBD, and their conformations control the activation of the machine and DnaK binding (12, 14-16). Mutants of ClpB with tilted MDs were found to be toxic *in vivo* (14, 17), highlighting the importance of tight control of the machine. We recently reported, based on single-molecule FRET (smFRET) spectroscopic studies, that the MDs of ClpB are highly dynamic, moving on the sub-millisecond timescale between their collinear (inactive) and tilted (active) states (18). We further showed that the MD is a continuous (analog), rather than a two-state (digital), activation switch for ClpB, with the population ratio of its states affecting the overall activity of ClpB. This population ratio was shown to be modulated by a range of allosteric signals, such as the binding of DnaK, nucleotides and substrate proteins. Although the dynamic detachment of the MDs is important, the full activation of ClpB also requires substrate-protein binding as an additional signal (12), suggesting a crucial involvement of its substrate-binding regions.

The NTDs of ClpB are 135-147-residue domains that form a distinct large ring at the top of the hexamer (Fig. 1b) (6). These domains, which are also conserved in the orthologs ClpA and ClpC, serve ClpB as initial binding sites for certain substrate-proteins (19, 20) prior to their processing. Because of its high mobility, the entire upper ring is frequently unobserved in cryo-electron microscopy (cryo-EM) reconstructions of ClpB hexamers (13, 15, 21, 22). However, two recent cryo-EM studies employed “focused classification” to capture several substrate-bound NTDs (23, 24). They revealed NTD trimers that made contacts with substrate protein molecules and surrounded the central channel of ClpB. In addition, a full ring of six NTDs was resolved in a cryo-EM structure of the yeast homolog of ClpB, Hsp104, also showing that they interact with the substrate (25).

Due to the difficulty in characterization, the experimentally validated functional role of the NTDs in ClpB remains limited to substrate-protein engagement. Their substrate-binding properties were firmly established by biochemical studies (19, 26, 27) and by cryo-EM (23-25), and the substrate-binding groove was characterized in high detail by NMR spectroscopy (20). However, accumulating experimental evidence suggests that their role extends beyond substrate-protein binding. Numerous studies reported increased ATP hydrolysis rate by the NTD-truncated mutant ΔNClpB in the absence of protein substrates, compared to the full-length ClpB, indicating that the NTDs may inhibit futile ATP hydrolysis through yet uncharacterized communication pathways (10, 19, 20, 28-33). Indeed, potentially relevant interactions of the NTD with the MD and NBDs in solution were recently detected by X-ray footprinting in Hsp104 (34). Furthermore, the function of the NTDs during substrate processing and translocation remains uncertain. Recently, an NTD deletion was reported to abolish the translocation of maltose-binding protein (35). On the other hand, ΔNClpB remained active in multiple *in vitro* disaggregation assays in the presence of DnaK-DnaJ-GrpE with other pre-aggregated protein substrates, implying a secondary role for the NTD in this process (20, 26, 27, 29, 36). It was proposed that the NTDs block the central channel (20) and can undergo conformational changes to actively assist with the translocation and disaggregation of substrate-proteins (27). NMR spectroscopy in aqueous solution suggested fast movement of the NTDs (20). These dynamics have not been studied in detail and the timescales of the motion, any potential conformational rearrangements and their implications for the disaggregase function remain to be characterized.

Here, we used a combination of single-molecule FRET (smFRET) spectroscopy and biochemical experiments to shed new light on the roles of the NTDs in ClpB. We directly monitor the sub-millisecond motions of the NTDs and demonstrate that engagement of substrate proteins does not restrict these conformational dynamics. We characterize the communication of the NTD with other functional domains of ClpB. Our data identify two independent allosteric pathways that stem from the NTD and are both important for the control of ClpB’s activation. Direct contacts of the NTD residues with the MD modulate its conformation, maintaining the inactive state of the MD. In addition, residues within the flexible linker connecting the NTD to NBD1 suppress the ATPase and disaggregation activities of ClpB. These results suggest that the NTD is critically involved in the reversible activation of ClpB through tight coupling to other functional domains, and is engaged in at least two independent allosteric communication pathways through its ultrafast motions.

## Results

### The NTD fluctuates on the microsecond timescale with minor changes in conformational dynamics upon substrate binding

In the crystal structure of the full-length *Thermus thermophilus (TT)* ClpB (henceforth ClpB) (6), the NTDs are connected to the neighboring NBD1 domains by disordered linkers, which might suggest that these domains are highly mobile. Analysis of the NTD dynamics in a related AAA+ hexameric machine p97 by solution NMR spectroscopy suggested microsecond motions (38). Here, we set to characterize the real-time dynamics of the NTD within the ClpB hexamer in aqueous solution. As in our recent study of the MD dynamics in the full-length ClpB (18), we employed a powerful combination of smFRET and photon-by-photon hidden Markov model analysis (H^2^MM), described in detail previously (39). To study the NTD dynamics, we located a cysteine for fluorescent labeling on the NTD, at the end of α-helix A1 (residue R23), and a second cysteine was inserted into a rigid position within NBD1 (residue D176) of ClpB (Fig. 1c). The resulting double mutant of ClpB was labeled with Alexa Fluor 488 (AF488) and Alexa Fluor 594 (AF594) (Methods). It was confirmed that the fluorescently-labeled 23C-176C construct exhibited unaltered ATP activity (3.2±0.1 min^-1^ relative to 3.5±0.2 min^-1^ in WT ClpB). It showed prominent ATPase stimulation upon binding of the model protein substrate κ-casein (40) (3.1-fold enhancement, Fig. 2a) and had no assembly defects (Fig. S1). Furthermore, fluorescence anisotropy measurements of single-cysteine mutants (23C and 176C) indicated that the dyes attached at these positions showed unrestricted rotation both in the absence and in the presence of κ-casein (Fig. S1). Double-labeled ClpB subunits were mixed with a 100-fold molar excess of unmodified unlabeled ClpB subunits (WT ClpB), ensuring that the newly assembled ClpB hexamers contained only one fluorescently labeled protomer.

**Figure 2.**
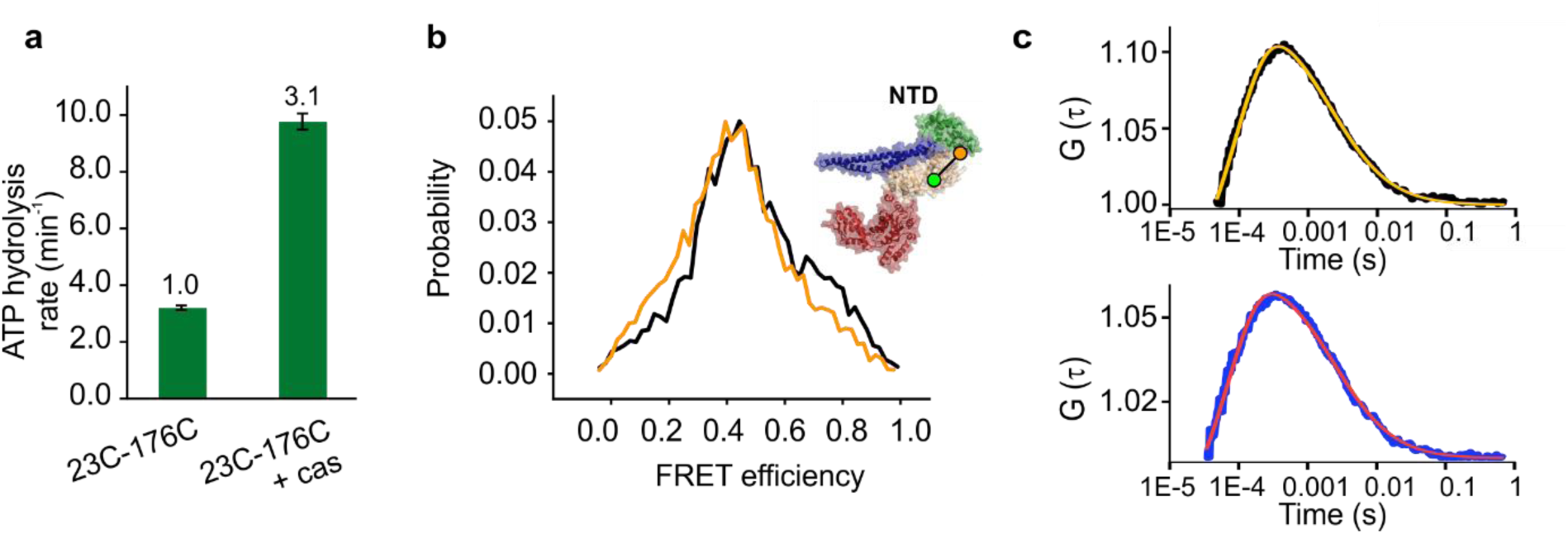
Analysis of the conformational dynamics of the NTD. **(a)** ATPase activity of fully-labeled 23C-176C protein, either basal or in the presence of 25 µM κ-casein (n=3). The error bars correspond to standard errors of the mean. The numbers above the bars designate relative activity. **(b)** FRET efficiency histograms of the 23C-176C mutant of ClpB (2 mM ATP) in the absence of substrate-protein (black) and in the presence of 25 µM κ-casein (yellow). Here and elsewhere, FRET efficiency histograms are area-normalized. **(c)** Filtered FLCS cross-correlation curves of 23C-176C ClpB without (black, top) and with 25 µM κ-casein (blue, bottom). The yellow and red curves are fits to equation 1 in Methods and yield similar diffusion coefficients but faster dynamics in the presence of κ-casein - see text.

We conducted photon-by-photon smFRET measurements on diffusing molecules of the construct introduced above in the presence of 2 mM ATP. The arrival times of the photons emitted by both donor (AF488) and acceptor (AF594) dyes were registered on two separate detectors. After selecting computationally molecules that contained both AF488 and AF594 dyes (Methods and Fig. S4), we calculated FRET efficiency histograms. The histograms were broad (Fig. 2b), indicating multiple conformations of the NTD. Surprisingly, there was only a minor change in the appearance of the histograms measured in the presence of κ-casein, even while previous studies proposed that the NTDs direct bound substrate-proteins towards the central pore (27).

To assess the timescale of the NTD motion within ClpB hexamers, we analyzed the dynamics by fluorescence lifetime correlation spectroscopy (FLCS). This experimental technique (41) enables filtering fluorescence cross-correlation functions to obtain the contribution of separate sub-species observed in the FRET histogram (see Methods). We calculated cross-correlation functions between the species with the lowest FRET efficiency values (<0.18) and the species with the highest FRET efficiency values (>0.75). The resulting curves (Fig. 2c) showed a prominent rise of the signal on the microsecond timescale, and analysis indicated conformational dynamics with an exchange rate of 8,330±630 s^−1^. Interestingly, the addition of κ-casein at a concentration of 25 µM led to acceleration of the dynamics to 12,500±780 s^−1^.

We also analyzed the NTD dynamics by applying H^2^MM (39), using a model with three states, which was the smallest number of states required to properly fit the smFRET data (see below). We did not attempt to assign these three states, but rather used them to derive dynamic information and compre to the FLCS results. The transition rates out of states 1, 2 and 3 were 3,540±480 s^−1^, 1860±150 s^−1^ and 2500±220 s^−1^, respectively, and were increased in the presence of κ-casein (Table S1), consistent with the FLCS analysis. To validate these results, we performed analysis of dwell time distributions (18, 42), and the resulting transition rates were in close agreement with the values obtained from H^2^MM (Table S1), suggesting that three states fit the data well. From the H^2^MM-derived FRET efficiency values of states 1 and 3 (0.18 and 0.81), we calculated the amplitude of the NTD motion to be ∼28 Å. Together, these results show that, within the hexamer, the NTD undergoes ultrafast large-scale movements that are independent of nucleotide state and are only mildly accelerated by the binding of the model substrate-protein. This high speed and conformational freedom are likely to underlie all of the NTD-driven regulations in ClpB that we characterize in subsequent sections.

### The NTD suppresses ATPase and disaggregation activities of ClpB

To further understand the role of the NTD in ClpB’s activity, we expressed a truncated version, ΔNClpB, which lacks the first 140 residues of the protein (Fig. S3). We confirmed by native polyacrylamide gel electrophoresis that both ClpB and ΔNClpB were well-assembled (Fig. S2). Measurements of ATP hydrolysis rates in the presence of 2 mM ATP at 25°C yielded a doubled value for ΔNClpB compared to ClpB (6.8±0.3 min^-1^ vs. 3.5±0.2 min^-1^) (Fig. 3a), in close agreement with previous literature values (18, 29, 43). However, in the presence of κ-casein (50 µM), similar maximal hydrolysis rates were reached by ClpB and ΔNClpB (10.4±0.4 min^-1^ and 9.6±0.6 min^-1^, respectively, Fig. 3a). The results suggest that the truncated ClpB variant is dysregulated and exhibits futile ATPase activity in the absence of protein substrate. We compared the disaggregation activity of ClpB and ΔNClpB by monitoring the reactivation of heat-induced aggregates of glucose-6-phosphate dehydrogenase (G6PDH) (44) and observed a ∼67% regeneration of soluble G6PDH activity by both ClpB and ΔNClpB (Fig. 3b). Even though the yields of disaggregation by the two variants were essentially the same, ΔNClpB displayed a higher disaggregation rate compared to the full-length variant (3.0±0.1 nM.min^-1^ vs. 2.2±0.1 nM.min^-1^, Fig. 3c). This large difference in the rates of disaggregation was also observed in experiments with heat-induced aggregates of firefly luciferase (21.7±0.5 min^-1^ for ΔNClpB compared to 13.1±0.65 min^-1^ for ClpB) (Fig. 3d).

**Figure 3.**
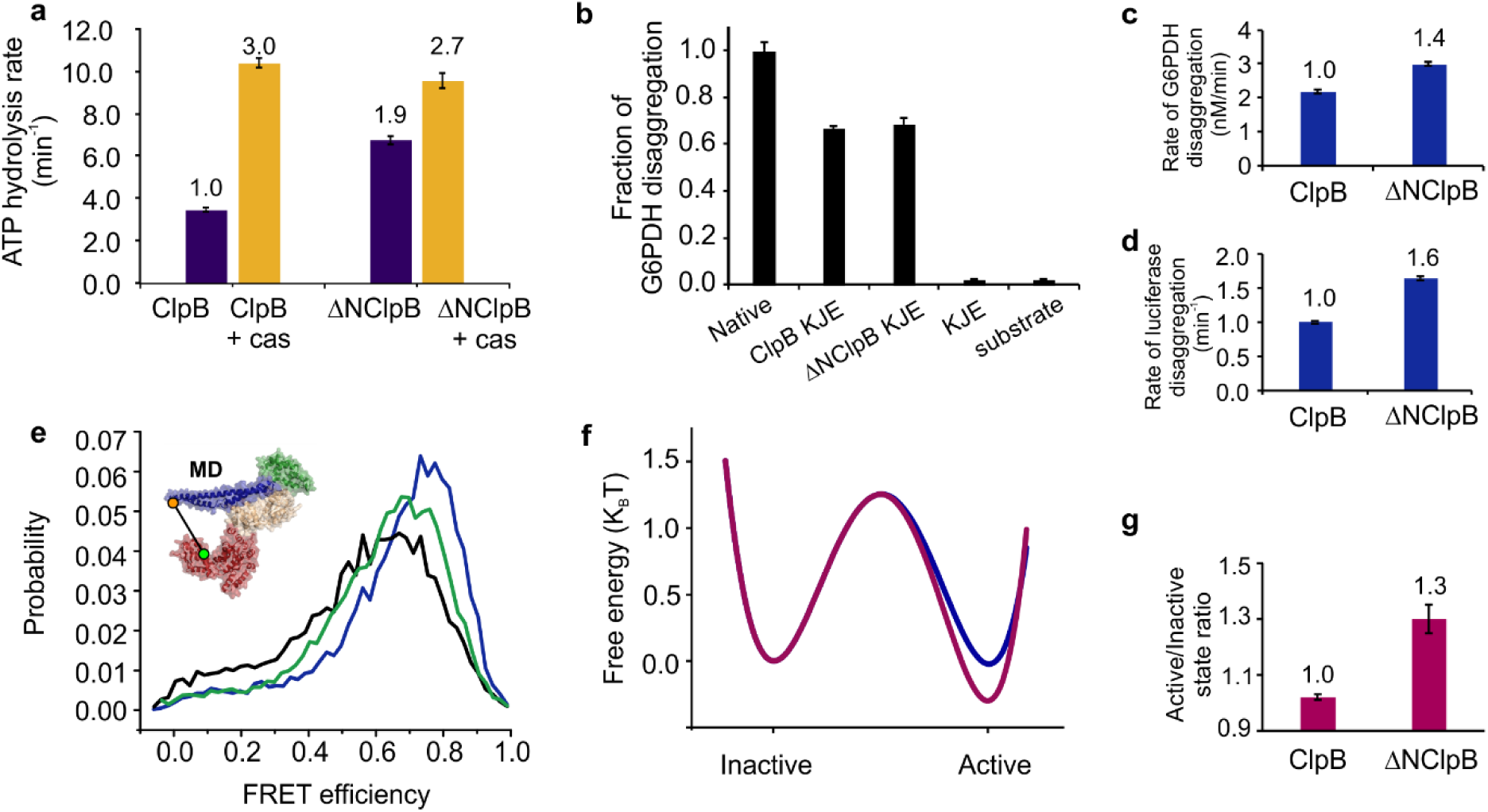
ATPase, disaggregation activities and MD dynamics of ClpB and ΔNClpB. **(a)** Rate of ATP hydrolysis measured at 25°C with and without 50 µM κ-casein (“cas”, n=3). **(b)** Disaggregation of heat-induced aggregates of G6PDH as a substrate, measured after 3 h of incubation with ClpB variants (2 µM) in the presence of DnaK, DnaJ and GrpE (KJE, 2 µM, 1 µM and 1 µM, respectively). The fraction of disaggregation was determined by taking the activity of native (non-aggregated) G6PDH as 1 (n=6). **(c)** Rate of disaggregation of heat-induced aggregates of G6PDH or of firefly luciferase **(d)** by ClpB variants in the presence of DnaK, DnaJ and GrpE (n=3). **(e)** FRET efficiency histograms of the full-length S428C-S771C construct of ClpB (black) and of the respective ΔNClpB mutant in the presence of 2 mM ATP (green) and in the presence of 25 µM κ-casein (blue). **(f)** Comparison of the free-energy profiles of MD dynamics of the full-length ClpB (blue) and ΔNClpB (purple), calculated from H^2^MM parameters using the Arrhenius equation with a pre-exponential factor of 10^5^ s^−1^. **(g)** The population ratio of horizontal (inactive) to tilted (active) states of the MD, derived from H^2^MM analysis. The error bars in all panels correspond to standard errors of the mean.

### smFRET experiments reveal that ΔNClpB has activated MDs

To shed light on the effect of the NTD deletion on ClpB activity, we probed the deletion influence on MD dynamics by using smFRET measurements and H^2^MM analysis (18, 39). In our recent study on the MD dynamics of ClpB in aqueous solution (18), single-molecule trajectories were fitted with a tree-state model, and the validity of the selected model was unequivocally validated through multiple additional methods. In that study, we used measurements with three separate FRET pairs to assign the two major (most populated) states of the MD, state 1 and 2, to the active (tilted) and inactive (collinear) conformations of the MD, respectively. Strikingly, in the absence of substrates or co-chaperones, the two states in the full-length ClpB were populated equally, with an active/inactive population ratio of 1.00 ± 0.01, and were rapidly interconverting with transition rates of *k*_12_ = 5300±150 s^−1^ and *k*_21_ = 5700±100 s^−1^, respectively.

To characterize the MD dynamics of ΔNClpB by smFRET, we generated a double-cysteine mutant of ΔNClpB (Fig. 1c). The first cysteine was introduced into motif 1 of the MD (residue 288, which corresponds to 428 in the full-length ClpB) and the second into NBD2 (residue 631, corresponding to 771 in the full-length ClpB, Fig. 1c). The resulting double-mutant of ΔNClpB was labeled with AF488 and AF594 (Methods), and mixed with a 100-fold molar excess of unmodified ΔNClpB to obtain a single fluorescently labeled protomer within each hexamer. We tested the effect of labeling on ΔNClpB, and found no change in ATPase or disaggregation activity (Fig. S2), in agreement with previous findings on the full-length ClpB (18). smFRET measurements were carried out on ΔNClpB in solution, in the presence of 2 mM ATP. The addition of ATP guaranteed the stability of ClpB hexamers in all our smFRET experiments; indeed, FRET efficiency histograms showed no evidence for their dissociation during the measurements (Fig. S5).

FRET efficiency histograms of ΔNClpB appeared to be shifted to higher average FRET efficiency values compared to the data for the full-length ClpB (Fig. 3e). The two major states of the MD were found to possess FRET efficiency values of 0.8±0.01 (state 1) and 0.48±0.01 (state 2), in good agreement with the active and inactive states of the full-length variant (18). In contrast to the results for the full-length ClpB, though, the states were unequally populated, with relative populations of 0.56±0.02 and 0.44±0.02, respectively, and an increased active/inactive state ratio of 1.30±0.05 instead of 1.00±0.01 (Fig. 3g), commensurate with the overall shift of the FRET histogram to higher value. As in the full-length ClpB, the two states were found to be under fast exchange, with interconversion rates of *k*_12_ = 4800±200 s^−1^ (from active to inactive state) and *k*_21_ = 6300±90 s^−1^ (reverse direction) (Table S3). Approximate free energy profiles of the MD dynamics, constructed based on the derived rates, showed that the energy barrier for the transition from inactive to active state was decreased in ΔNClpB, and the active state was stabilized (Fig. 3f). These results indicate that the MD in ΔNClpB variant toggles between the same conformations as in the full-length variant, but, remarkably, favors the active conformation. Based on our previous findings, the higher active/inactive state ratio should lead to a larger disaggregation rate of ΔNClpB (18), as is indeed observed (Fig. 3 c,d). To find whether or not this activation of the MD in ΔNClpB could be enhanced further by protein substrates, we repeated the experiments in the presence of 25 µM κ-casein. The resulting FRET efficiency histograms showed higher average FRET efficiency values (Fig. 3e, Tables S2 and S3) due to further increase in the active/inactive state ratio to 1.76±0.01, indicating that additional activation of the MD in ΔNClpB can take place in response to substrate binding.

### The NTD suppresses the MD through direct contacts

Next, we set out to explain why the MD appears activated upon the deletion of the NTD. It was previously found biochemically that the isolated NTD of ClpB from *E. coli* does not form stable contacts with other parts of the molecule (45), and we confirmed this result for *TT* ClpB, used in this study. ΔNClpB and the isolated NTD, comprising residues 1-141 of ClpB, that were mixed together (at 40 µM ΔNClpB and 650 µM NTD) migrated separately in size exclusion chromatography and native polyacrylamide gel electrophoresis analysis (Fig. S6), indicating that no stable complex was formed.

Since the NTD is connected to the neighboring NBD1 by a flexible linker region (6), it is interesting to ask whether its rapid motions in aqueous solution allow it to reach the MD. We first qualitatively assessed this possibility by using a simple computational procedure that sampled the conformational space of the NTD (see Methods section). From the inspection of the crystal structure of ClpB (6), we found that the region of the MD that is closest to the NTD is the edge of motif 2. Starting with the crystal structure of the ClpB monomer (6), we generated multiple conformations of the NTD within the context of the full hexamer by fully rotating it around a single residue in the flexible linker region (position 142). Then, we selected NTD residues that were located within 10 Å from the edge of motif 2 (residue 487) in the generated conformations (Fig. 4a), and found 11 in total, 5 of which were on or close to α-helix A1 of the NTD (these residues are indicated in Fig. S3). Based on this finding, we designed ClpB mutants in order to test whether we can experimentally detect an interaction between α-helix A1 and the MD. To this end, we used Atto 655, a fluorophore that is effectively quenched upon the interaction with several amino acids, especially with tryptophan (W), leading to a significant decrease in fluorescence signal (46). We introduced a cysteine into the MD (position 487) of the full-length ClpB for labeling with maleimide-linked Atto 655. We then incorporated single tryptophan residues into the NTD, either at position 12 or 23 within α-helix A1 (Fig. S3), and used a mutant without these additional tryptophans as a control. Fully-labeled ClpB mutants Atto 655-487C-ClpB, Atto 655-487C-12W-ClpB and Atto 655-487C-23W-ClpB were found to be well-assembled in the presence of 2 mM ATP and showed ATPase activity that was comparable to that of unmodified ClpB. 487C-ClpB variants were fully active in disaggregation experiments before labeling, but had no disaggregation activity when fully-labeled, likely due to the hindrance of the DnaK binding site (16) on the MD by Atto 655 (Fig. S7).

**Figure 4.**
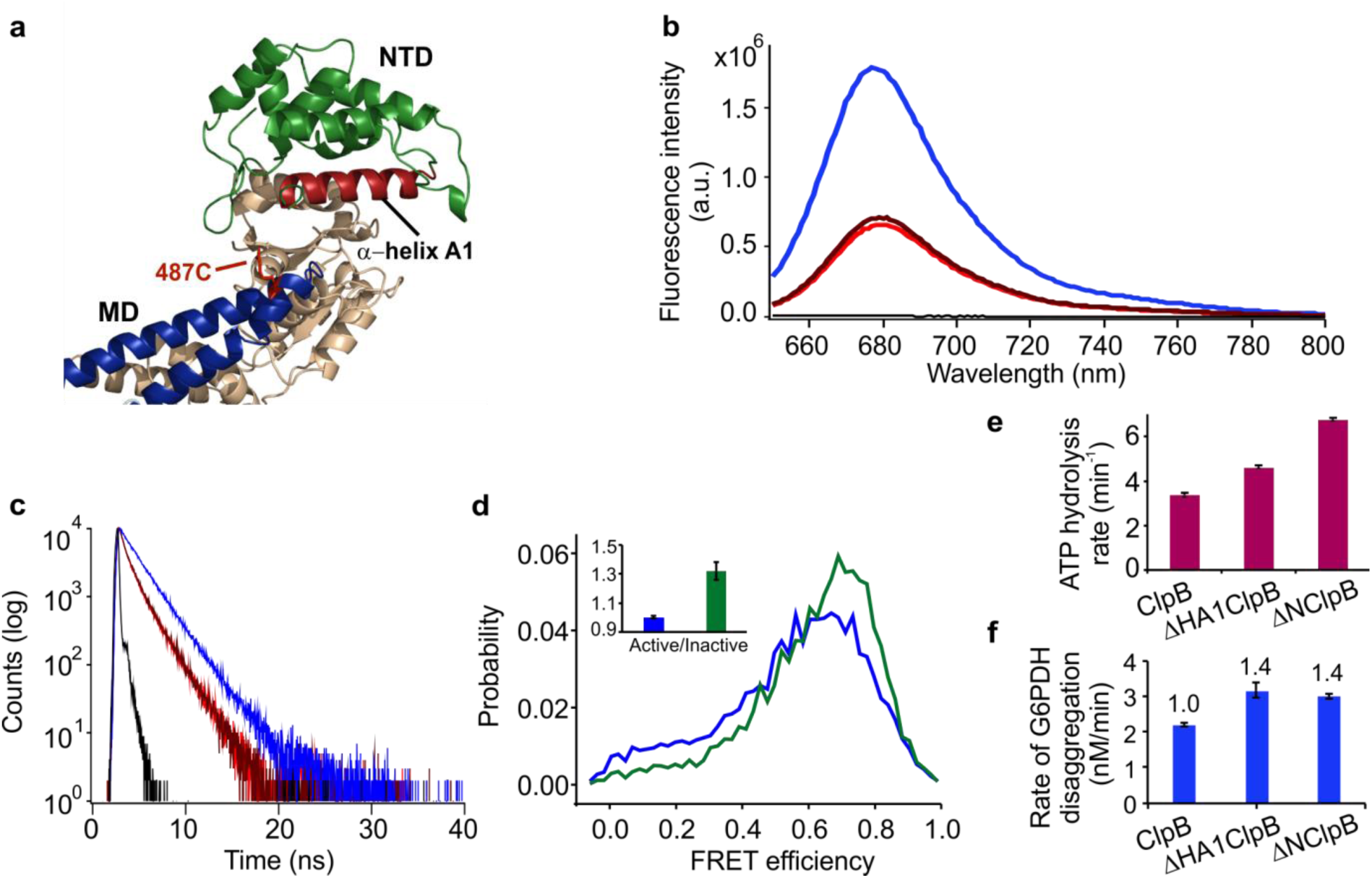
NTD mutants of ClpB affect MD dynamics. **(a)** Zoom into the upper region of ClpB protomer (crystal structure PDB:1QVR) (6). The MD is colored in blue, and the NTD in green, with α-helix A1 of the NTD highlighted in red. Residue 487C on the MD used for Atto 655 incorporation is shown as red sticks. **(b)** Steady-state fluorescence spectra of Atto 655-labeled ClpB mutants: ClpB (blue), 12W-ClpB (red), 23W-ClpB (dark red), buffer (black). Average fractions of fluorescence intensity, normalized to the result of the mutant without incorporated tryptophan, were 0.36 for 12W and 0.39 for 23W. **(c)** Fluorescence decay curves: ClpB (blue), 12W-ClpB (red), 23W-ClpB (dark-red), instrument response function (black). The resulting fluorescence lifetimes were 2 ns for the control ClpB mutant, and 1.3 ns for 12W and 23W variants. **(d)** FRET efficiency histograms of the full-length ClpB (blue) and the truncated ΔHA1ClpB variant (green). Inset shows the corresponding active/inactive state ratios of the MD. **(e)** ATPase activity of ΔHA1ClpB, compared to the full-length and ΔNClpB variants (n=4). **(f)** Rate of disaggregation of G6PDH aggregates in the presence of DnaK, DnaJ and GrpE (n=3). The error bars correspond to standard errors of the mean.

Bulk steady-state fluorescence measurements of the Atto 655-labeled variants (Fig. 4b) showed a strong decrease (around 60%) of the fluorescence intensity in the tryptophan-containing mutants relative to the control 487C-ClpB, and the fluorescence lifetimes of these variants were found to be shortened (from 2 ns in the control samples to 1.3 ns in 12W- and 23W-containing variants). These differences indicate quenching of the fluorophore by the tryptophan residues in the NTD. Since the fractional decrease in fluorescence intensity of the mutants was larger than the decrease in fluorescence lifetime, we concluded that the quenching process has both a static component, due to the formation of a non-fluorescent Atto 655-quencher complex, and a dynamic component, due to transient encounters of the fluorophore with the quencher (47). This finding is commensurate with previous results on the effect of W on Atto 655 fluorescence (46). Considering that the dye is located on the tip of the MD, these results are in agreement with our hypothesis that the NTD can make contacts with the MD.

In order to determine whether the contacts that the NTD makes with the MD are sufficient to affect the MD dynamics, we removed the entire α-helix A1 from the NTD by deleting residues 8-25 of the full-length ClpB (Fig. 4a, Fig. S3). This truncated mutant (ΔHA1ClpB) was fully assembled (Fig. S7), and smFRET analysis of its double-labeled variant revealed that its MD was activated in comparison with the MD of the full-length ClpB. Indeed, FRET efficiency histograms (Fig. 4d) were shifted to higher FRET values when compared with the histograms for the full-length ClpB. The derived interconversion rates of the MD were unequal, *k*_12_ = 4400±100 s^−1^ and *k*_21_ = 5900±300 s^−1^ (Table S3), resulting in an increased active/inactive state ratio of 1.32±0.06, comparable to the ratio found in ΔNClpB (Table S2). Measurements of the ATP hydrolysis rate of ΔHA1ClpB yielded a value of 4.6±0.1 min^-1^ (Fig. 4e), intermediate between the activity of the full-length ClpB (3.5 min^-1^) and ΔNClpB (6.8 min^-1^). The rate of disaggregation of heat-treated G6PDH by ΔHA1ClpB was higher than by the full-length ClpB (3.2±0.2 nM.min^-1^ vs. 2.2±0.1 nM.min^-1^, Fig. 4f). Therefore, removal of the interactions with α-helix A1 of the NTD activates the MD. This is consistent with the fluorescence quenching results, which indicate a direct contact of α-helix A1 with the MD. These results imply that α-helix A1 comprises residues that can reach the MD and significantly affect its conformational transitions.

### The NTD further affects the activity of ClpB through an MD-independent pathway

Given that the interaction of the α-helix A1 with the MD could not account for the full effect of the NTD on ClpB functional measures, we looked for additional potential regions of the NTD involved in interaction with the rest of the protein. To this end, we investigated a well-conserved region within the flexible linker that connects the NTD to NBD1 and is absent in ΔNClpB (6). Flexible interdomain linkers are known to be involved in allosteric regulation in multiple proteins (48), especially through charged residues. We substituted a conserved charged residue R136 for an uncharged glycine. The resulting ClpB mutant R136G was found to be well-assembled (Fig. S8), displayed an increased ATPase rate, 6.7±0.1 min^-1^, matching the value obtained for ΔNClpB under our experimental conditions (6.8 min^-1^) (Fig. 5a), and showed a higher G6PDH disaggregation rate (3.9±0.2 nM.min^-1^ vs. 2.2±0.1 nM.min^-1^ for unmodified ClpB, Fig. 5b). To test whether this effect was specific to R136, we separately mutated neighboring residues, 134 or 135, to glycine (Fig. S3). The ATPase activity of these single mutants was not raised above the value for unmodified ClpB (3.7±0.3 min^-1^ and 3.5±0.4 min^-1^ for E134G ClpB and L135G ClpB, respectively) (Fig. 5a) and the rates of G6PDH disaggregation were unaltered (Fig. 5b), confirming that R136 is selectively involved in ClpB regulation.

**Figure 5.**
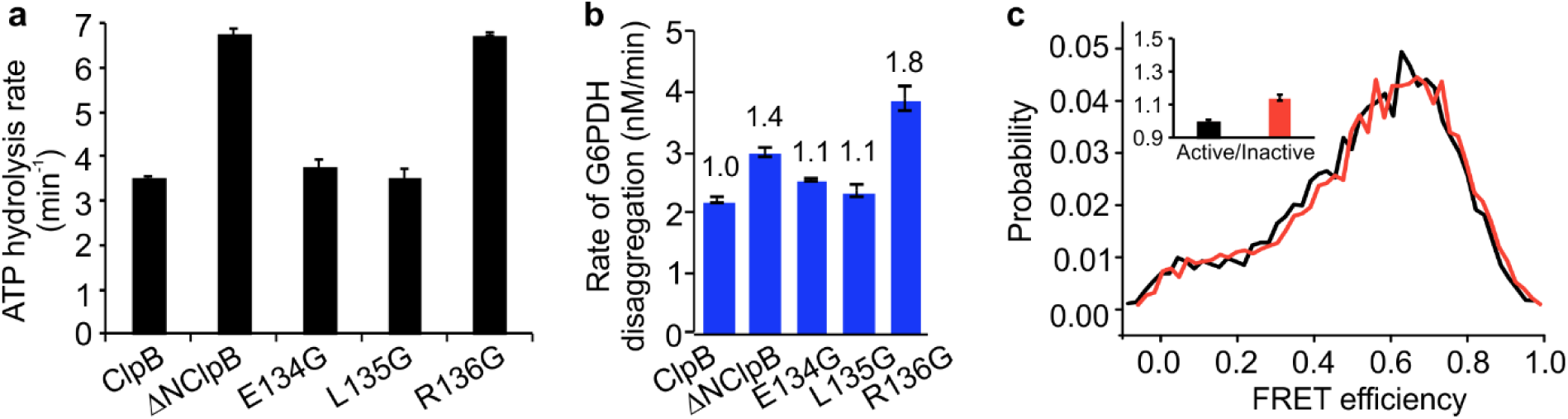
Residue R136 regulates ATPase and disaggregation activities but does not affect MD dynamics. **(a)** Comparison of ATP activity of the full-length ClpB and ΔNClpB with point mutants in the flexible linker region, residues 134-136 (n=4). **(b)** Rate of disaggregation of G6PDH aggregates in the presence of DnaK, DnaJ and GrpE (n=2-4). **(c)** FRET efficiency histogram of the full-length R136G mutant (orange) compared to WT ClpB (black). Inset shows the active/inactive state ratios of the MD. The error bars correspond to standard errors of the mean.

To determine whether the MD dynamics were affected by this pathway, we generated R136G ClpB molecules with a single double-labeled subunit (S428C-S771C), and measured them by smFRET. The FRET efficiency histogram of this mutant was similar to that of WT ClpB (Fig. 5c). H^2^MM analysis yielded a value of 1.14±0.02 for the active/inactive state ratio of the MD, which is only slightly increased over 1.00±0.01 in the unmodified ClpB (Table S2). Therefore, there seems to be no major effect on the MD population ratio following R136G substitution, indicating that the NTD-NBD communication pathway that is responsible for the increased ATPase and disaggregation rates in the R136G variant of ClpB does not involve the MD.

Because R136G is a charged residue, it might form salt bridges that underlie its role in the ATPase regulation. Interestingly, a previous biochemical study reported the formation of an intra-subunit disulfide bond in a R136C-Q260C mutant of *TT* ClpB (27), strongly suggesting that in solution, R136G can reach Q260 residue. The latter is located within NBD1 in close proximity to a conserved Walker B region (aas 266-271), which is responsible for ATP hydrolysis (49). Therefore, it is plausible that R136G forms a salt bridge with a residue within NBD1 (E257 or E264), close to Walker B region. We will test this conjecture in future work.

### NTD removal does not disrupt communication between the MD and NBDs

Several mutations that are located remotely from the NTD or the NTD-NBD1 linker have been shown to activate or repress the activity of ClpB (14, 15, 17). We asked whether these mutations still have the same effect in ClpB, after its NTD is deleted. The MD is known to be tightly regulated through a network of salt bridges that connect it to the neighboring NBD1 (14, 15, 17). To check whether this regulatory pathway is retained upon the NTD deletion, we generated activated and repressed hexamers of ΔNClpB by introducing previously characterized single mutations that were expected to alter the MD dynamics (Fig S3). Point-mutation E209A in NBD1 (termed hyper-ΔNClpB, corresponding to E349A in the full-length ClpB) breaks a salt bridge connecting it to NBD1, which results in a detached and activated MD (17). The hyper-ΔNClpB variant showed 3.5-fold higher rate of ATP hydrolysis than the unmodified ΔNClpB, and the same disaggregation yield as ΔNClpB (Fig. 6a, b). These results are consistent with the reported effects of similar mutations in the full-length ClpB (18) and in ΔNClpB from *E. coli* (50). The FRET efficiency histogram of the hyper-ΔNClpB variant was shifted to higher values compared to ΔNClpB data (Fig. 6c). The transition rates of the MD, extracted from the analysis of smFRET trajectories of this mutant were altered (Table S3), *k*_12_ = 4100±400 s^−1^ and *k*_21_ = 7800±400 s^−1^, resulting in the relative populations of the active and inactive states of 0.65±0.01 and 0.35±0.01. Consequently, the active/inactive state ratio was 1.85±0.07 (Table S2), higher than in ΔNClpB, and similar to the result for the activated mutant of the full-length protein (18).

**Figure 6.**
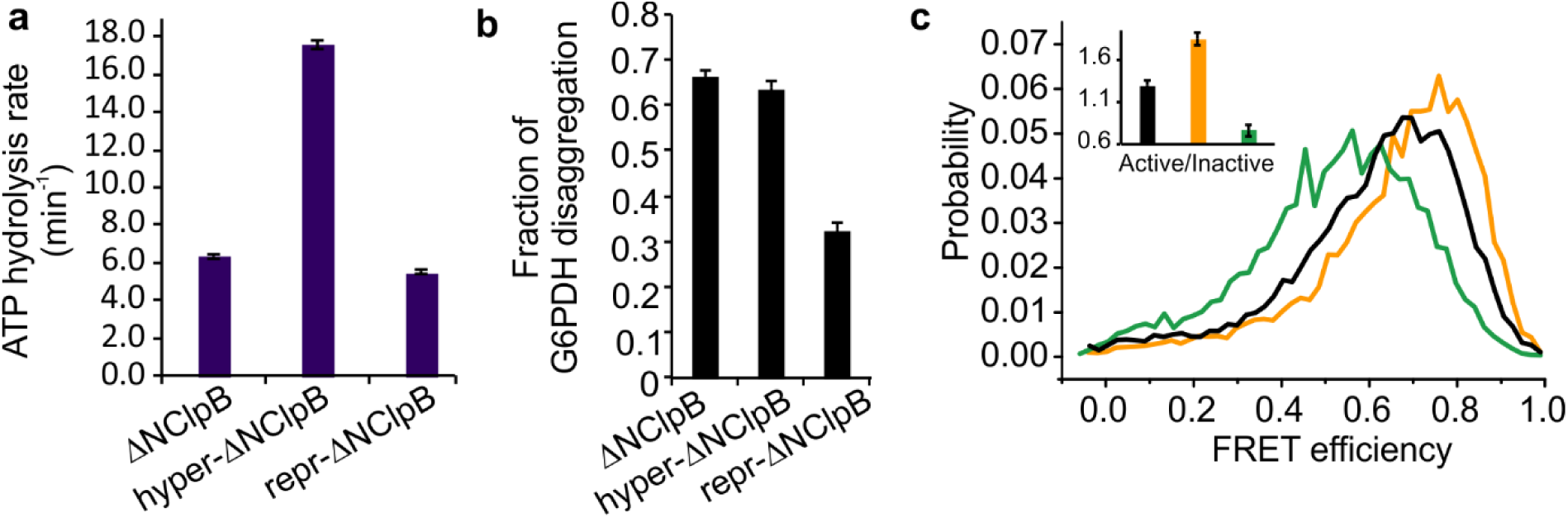
Characterization of MD-NBDs allosteric pathways in ΔNClpB. **(a)** The rate of ATP hydrolysis of activated and repressed ΔNClpB mutants at 25°C (n=3-5). **(b)** Disaggregation of heat-induced aggregates of G6PDH by ΔNClpB mutants, measured after 3 h disaggregation in the presence of DnaK, DnaJ and GrpE (n=2). **(c)** FRET efficiency histograms of activated and repressed mutants of ΔNClpB: ΔNClpB (black), hyper-ClpB (yellow), repr-ΔNClpB (green). Inset shows the corresponding active/inactive state ratios of the MD. The error bars correspond to standard errors of the mean.

Point-mutation E283A within the MD (repr-ΔNClpB, corresponding to E423A in the full-length ClpB) stabilizes its contacts with NBD1 and with MDs of neighboring subunits (14, 15, 51, 52). Repr-ΔNClpB had slightly decreased (by 10%) ATPase activity compared to ΔNClpB, and its disaggregation activity was reduced by half compared to the non-mutated ΔNClpB (Fig. 6a,b), in good agreement with the previously observed effects of this mutation in the full-length protein (18, 51, 52). The FRET efficiency histogram of the repr-ΔNClpB was shifted to lower average values in comparison to ΔNClpB. The derived interconversion rates were *k*_12_ = 5500±150 s^−1^ and *k*_21_ = 4200±300 s^−1^ (Table S3), leading to an increased relative population of the inactive state (0.57±0.04) and the active/inactive state ratio of 0.76±0.07, in agreement with the result of this mutation in the full-length ClpB (Table S2) (18). Therefore, both the activating and repressing mutations strongly affected the MD dynamics of ΔNClpB. This suggests that the salt bridges between the MD and NBD1 that regulate the MD conformations in ClpB remain unaffected by NTD deletion.

We also probed another previously characterized pathway of MD regulation that involves the nucleotide binding sites of ClpB. Recently, we found that ATP binding to NBD1 decreased the population of the active state of the MD, whereas ATP binding to NBD2 had the opposite effect (18). We set to determine whether this regulation was retained in ΔNClpB. To this end, we generated and analyzed mutants of the Walker A motifs, which are located in the two NBDs and are responsible for ATP binding (Fig. S9) (49). We found no difference in their activity or in their MD dynamics in comparison to the corresponding full-length ClpB mutants (18), suggesting that NTD removal does not affect the allosteric regulation of the MD conformations upon nucleotide binding.

## Discussion

In this work, we studied the dynamics of the NTD and its communication with other domains of ClpB using a powerful combination of smFRET experiments, photon-by-photon H^2^MM analysis, fluorescence quenching measurements and biochemical assays. We found that the NTD undergoes fast motions and is involved in at least two separate allosteric routes of ClpB activation through the coupling to other functional domains.

Our smFRET analysis of the NTD dynamics in ClpB revealed that it moves on the microsecond timescale. This timescale should be contrasted with the expected relaxation time of a small domain like the NTD, which should be nanoseconds. This implies that some interactions, for example, salt bridges with NBD1, are limiting NTD dynamics. Indeed, from a closer inspection of the crystal structure of ClpB (*TT*) (6), the charged residues within α-helix A1 of the NTD can be found in close proximity to the oppositely charged residues within NBD1, comprising potential salt bridges E12-R228 and R23-E299. To note, the pair E12-R228 was previously mutated to cysteines for intra-molecular cross-linking, along with another already-mentioned pair R136-Q260 (27). An experimental investigation of the potential role of salt bridges in NTD dynamics is left to future work. Notably, our data suggest that the NTDs do not show major changes in their conformations relative to NBD1 upon substrate-protein binding. This is in stark contrast to p97, mentioned above, whose NTDs undergo a large upward movement with out-of-plane rotation of 75° upon activation (53). The absence of significant conformational changes in the NTD upon substrate-protein binding is in agreement with recent cryo-EM structure of ClpB, where a trimer of substrate-bound NBDs from alternating protomers was shown to form a ring at the channel entrance, maintaining the overall helical arrangement of the protomers (24). Nevertheless, the NTD moves faster by ∼50% upon substrate binding, which might have an effect on the dynamics of DnaK binding to the MD, thereby influencing the activation state of the whole machine.

Based on our results, ΔNClpB is an activated ClpB variant that has tilted MDs, elevated basal ATP hydrolysis rate and high disaggregation activity towards heat-induced aggregates of G6PDH and firefly luciferase. The increased basal ATP activity exhibited by ΔNClpB in comparison to the full-length variant is in good agreement with multiple preceding studies (10, 19, 20, 28-33), and implies that this truncated mutant of ClpB is more wasteful in terms of ATP consumption in the absence of bound substrate-proteins. Our data show unaltered yield and faster rate of disaggregation following the NTD removal. Thus, we can conclude that ΔNClpB is a more active disaggregase, at least towards certain protein substrates. The presence of a small fraction of the ΔNClpB variant, with its high intrinsic ATPase activity and increased disaggregation activity, might be crucial for cell survival under conditions of particularly harsh heat shock and stress.

Our data showed that α-helix A1 of the NTD makes geometric contacts with motif 2 of the regulatory MD of ClpB that significantly affect the dynamics of the latter. This interaction with the MD results in the suppression of ATPase activity and the inhibition of the disaggregation by ClpB (Fig. 7). This finding is unexpected: while the regulatory role of the MD was firmly established by multiple previous studies (12, 14-16, 18), the NTD is often viewed as a non-essential substrate-docking domain (29). The NTD-MD interaction identified here involves the functionally-important motif 2 of the MD. The detachment of motif 2 from the NBD1 of ClpB was shown to increase its ATPase activity (14). Moreover, this motif comprises the DnaK-binding epitope of ClpB (16), crucial to its disaggregation function (12, 15, 16). NTD-MD contacts were also inferred, though indirectly, from X-ray footprinting studies of the homologous Hsp104 (34). We directly verified the importance of the NTD-MD contacts by finding that the NTD removal resulted in tilting of the MD in ClpB, shifting its active/inactive population ratio from 1 to 1.3. Based on our experiments with activated, repressed and Walker A mutants of ΔNClpB, the effect is due to the loss of interacting residues within the NTD, rather than due to the disruption of any known MD regulations downstream of the NTD (18). Our data therefore suggest a yet unrecognized role for the NTD as a regulatory element that suppresses the MD of ClpB through direct contacts and controls its activation (Fig. 7).

**Figure 7.**
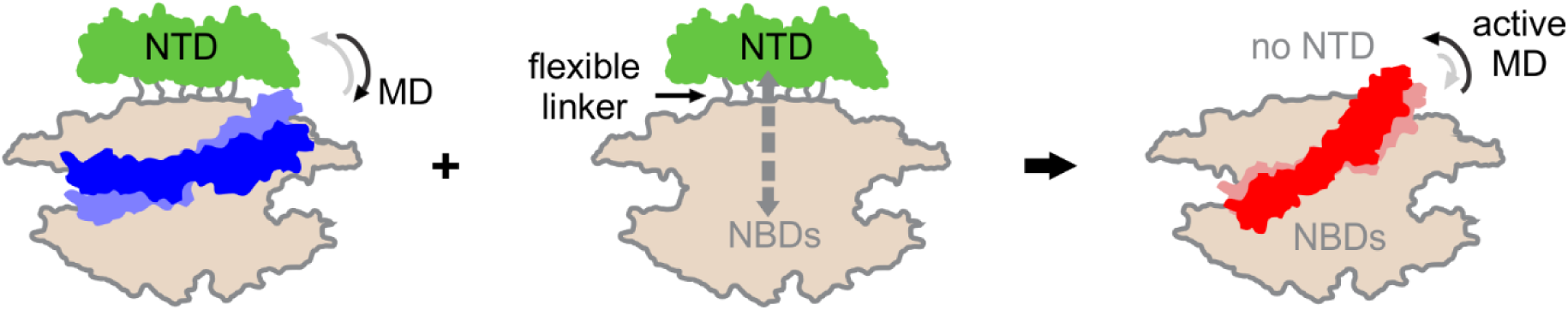
Schematic summary of the regulations by the NTD in ClpB. Left: In the full-length ClpB, the NTD (green) suppresses the MD (blue) through regulatory contacts. The MD is in dynamic equilibrium between active/inactive states with a state population ratio of 1. In addition, there is an MD-independent NTD-NBDs communication through the flexible interconnecting linker region. Both NTD-mediated regulations affect ATPase and disaggregation activities of ClpB. **Right:** In the ΔNClpB variant, which lacks the NTD, the MD suppression is lost, resulting in more populated active state of the MD (in red). The NTD-NBDs communication is absent. ΔNClpB exhibits higher ATPase and disaggregation activities due to the loss of both NTD-dependent regulations.

We found an allosteric residue R136 within the flexible linker region connecting the NTD to NBD1 that also regulates the ATPase and disaggregation activities of ClpB. The data showed, surprisingly, that mutation of this residue did not affect the MD conformations, yet increased the rate of ATP hydrolysis and accelerated the disaggregation of heat-induced aggregates of G6PDH. The effect of R136 mutation was specific since mutations of two preceding residues had no influence on these activities. To note, R136 is conserved in ClpB and Hsp104, although the sequence of the NTD-NBD1 linker region is highly variable (6). Given the proximity to NBD1, based on the X-ray structure of ClpB (6) and on previous biochemical findings (27), it is plausible that R136 residue forms a salt bridge that can affect the ATP-binding pocket to inhibit ATP hydrolysis, without any involvement of the surface-exposed MD. Indeed, the linker region within assembled hexamers of Hsp104 was reported to show a decrease in solvation by X-ray footprinting (34) and slowed hydrogen-deuterium exchange (54), suggesting that it becomes buried rather than surface-exposed.

## Conclusion

In this work, we characterized the microsecond-timescale dynamics of the substrate-binding NTD of ClpB, and found an unanticipated activity of this domain: it operates as an ultrafast control element for ClpB’s activity. As a result of the ultrafast motion, the NTD participates in two allosteric pathways: it suppresses the key regulatory MD through direct contacts, and also mediates allosteric signals to NBD1. These two pathways are totally independent of each other, and have functional implications, clearly displayed by the ΔNClpB mutant. The deletion of the NTDs causes the loss of contacts with the MDs and defective NTD-NBDs communication. As a consequence, ΔNClpB shows increased basal ATP activity and higher disaggregation rate than the full-length protein. We conclude that the NTD is involved both in substrate binding and in the regulation of machine activation. Thus, this domain is designed so that upon substrate-protein binding, it allosterically unlocks the inhibitory state of ClpB to optimally drive disaggregation. A similar multimodal allosteric control through a flexibly connected and fast-moving domain might be present in numerous multi-domain AAA+ molecular machines to enable their rapid adaptation to changing cellular conditions.

## Methods

### Protein expression and purification

*Thermus thermophilus* ClpB (*TT*. ClpB), its ΔNClpB variant beginning at residue 141 (Val), its mutants (Fig. S3), co-chaperones DnaK, DnaJ and GrpE, cloned into a pET28b vector, and the isolated N-terminal domain of ClpB (residues 1-141), cloned into pProEx vector, were expressed and purified according to recently published protocols (18), as detailed in the Supplementary Methods.

### ATP activity measurements

ATP activity of ClpB variants was measured using a coupled colorimetic assay (55). ClpB or its mutants (1 µM total monomer concentration) were incubated with 2 mM ATP and an ATP regeneration system (2.5 mM phosphoenol pyruvate, 10 units/ml pyruvate kinase, 15 units/ml lactate dehydrogenase, 2 mM 1,4 dithioerythritol, 2 mM EDTA, 0.25 mM NADH) in 50 mM HEPES (pH 8), 50 mM KCl and 0.01% Tween 20. For the experiments in the presence of the model substrate κ-casein (Sigma Aldrich), it was added to a final concentration of 50 µM. ATP hydrolysis was initiated by the addition of MgCl_2_ (10 mM), and measured by monitoring the time-dependent decrease in NADH absorption at 340 nm at 25°C. Data were background-corrected in all cases, and the measured ATP hydrolysis rate per ClpB monomer per minute is presented.

### Disaggregation of heat-induced aggregates of G6PDH and luciferase

Disaggregation experiments were performed following previously described procedures (18, 44, 56). The protocol for aggregate preparation and disaggregation reactions is detailed in the Supplementary Methods.

### Labeling of ClpB and ΔNClpB variants

Labeling of 23C-176C mutant of ClpB, S288C-S631C ΔNClpB and any of its mutational variants was performed similarly to the previously reported protocol (18). First, the double-cysteine mutant of ClpB (or ΔNClpB) was incubated for 1 h with 1:1.2 molar ratio of Alexa Fluor (AF) 594 (C5 maleimide, Invitrogen) under native conditions (25 mM HEPES, 25 mM KCl, pH 7). The unreacted dye was removed on a desalting column (Sephadex G25, GE Healthcare). The protein was exchanged into buffer containing guanidinium chloride in order to fully expose unlabeled cysteine residues (25 mM HEPES, 25 mM KCl, 2 M GdmCl, pH 7) and incubated with 1:1.5 molar ratio of AF488 (C5 maleimide, Invitrogen) for 1 h, following the separation of unreacted dye on a desalting column. Labeling was confirmed by absorption measurements. Labeling of the full-length 487C-ClpB variants with Atto 655-maleimide (Sigma Aldrich) was carried out by incubating ClpB with the dye at 1:1.5 ratio for 5 h in the dark (25 mM HEPES, 25 mM KCl, 2 M GdmCl, pH 7), followed by the separation of unreacted dye on a desalting column (Sephadex G25, GE Healthcare), buffer-exchange to native buffer (25 mM HEPES, 25 mM KCl, pH 7) and filtration of the labeled protein through 0.22 µm filter (Millex, Millipore).

### Mixing of fluorescently-labeled and unlabeled protomers

For the preparation of ClpB and ΔNClpB assemblies for smFRET experiments, AF488-AF594-labeled double-cysteine mutants were combined with 100-fold molar excess of unlabeled cysteine-less ClpB or ΔNClpB, respectively. This ratio ensured that the probability of the incorporation of one labeled protomer in a hexamer was 5.7%, whereas the probability to find two labeled protomers in the same hexamer was as low as 0.15%. To achieve full mixing, the protein solutions were initially dialyzed in the presence of 6 M GdmCl. This was followed by dialysis steps in the presence of 4 M, 2 M, 1 M and 0 M GdmCl. The final steps involved extensive dialysis into low-salt buffer (25 mM HEPES, 25 mM KCl, 10 mM MgCl_2_, 2 mM ATP, pH 8) and filtration through 0.1 µm filters (Whatman Anotop-10). The assembled ΔNClpB was aliquoted, flash-frozen and stored at −80°C until further use. For the preparation of activated, repressed ΔNClpB and its Walker A mutant hexamers, fluorescently double-labeled ΔNClpB bearing the respective mutation was mixed with the 100-fold excess of unlabeled ΔNClpB with the same mutation but without cysteine residues, and dialyzed according to the above protocol. For the preparation of R136G (or ΔHA1ClpB) hexamers of ClpB, double-labeled R136G (or ΔHA1ClpB) was mixed with the R136G (or ΔHA1ClpB, respectively) mutant of ClpB without cysteine residues.

### Single-molecule measurements

The preparation of custom-made glass flow chambers for single-molecule experiments was carried out following previously reported protocols (39, 42, 57). The chambers were coated with a supported lipid bilayer composed of egg phosphatidylcholine (Avanti Polar Lipids) to prevent protein absorption during measurements. The assembled hexamers of ClpB (or ΔNClpB) were diluted to ∼50 pM of labeled ClpB, corresponding to ∼5 nM of total ClpB, into single-molecule buffer (25 mM HEPES, 25 mM KCl, 10 mM MgCl_2_, 2 mM ATP, 0.01% Tween 20, pH 8). The solutions were loaded into the chambers, which were rapidly sealed to prevent evaporation. Measurements were conducted using a home-built inverted confocal single-molecule microscope described in detail previously (39, 42, 57). Data collection was carried out on freely diffusing molecules as reported (18). Briefly, the samples were illuminated with focused and overlapped 485 and 594 nm diode laser beams pulsed at a ratio of 3:1 with a repetition rate of 40 MHz. The emitted photons were divided into two channels using a dichroic mirror (FF580-FDi01, Semrock), and passed through band-pass filters, ET-535/70m for the AF488 emission and ET-645/75m for the AF594 emission (Chroma). The arrival times of the emitted photons in both channels were registered by single-photon avalanche photo-diodes (Perkin-Elmer SPCM-AQR-15) coupled to a standalone time-correlated single photon counting module (HydraHarp 400, PicoQuant). Data were acquired for up to 3.5 h per sample at ambient temperature (22°C). Between two and four individual samples were prepared and measured for each ClpB variant and condition.

### smFRET data analysis

Data analysis was performed as recently described (18). A cut-off of 10 µs was used to effectively separate fluorescence bursts from the background. FRET efficiency and stoichiometry were calculated as detailed elsewhere (58). We applied several corrections to eliminate any artifacts and to ensure that only fluorescent photons arising from individual molecules entered the analysis. More details on the data selection can be found in the Supplementary Information section, Fig. S4. FRET efficiency values were corrected for the leakage of photons from the donor to acceptor channel (estimated to be no more than 7%). Using stoichiometry/FRET efficiency histograms (Fig. S4), any singly-labeled species were eliminated and only the molecules containing both AF488 and AF594 were selected for further analysis. For the photon-by-photon H^2^MM analysis, fully detailed previously (39), the photons arising from pre-selected molecules after donor-only excitation were used. A fixed number of molecules (5,800) was analyzed for each sample. The data were fitted with a three-state model. In the case of the analysis of MD dynamics, relative populations of the two major states were extracted and compared between different ΔNClpB mutants or conditions (Table S2). This fitting approach was rigorously tested and confirmed through multiple additional methods on the equivalent full-length ClpB datasets in our preceding study (18).

### FLCS measurements

Conventional smFRET measurements of freely-diffusing double-labeled molecules of 23C-176C hexamers (assembled in 1:100 ratio with WT ClpB) were performed. Pulsed interleaved excitation by 485-nm and 594-nm lasers was used (in 1:1 sequence). The measurements were conducted with laser powers of 50 µW (485 nm) and 10 µW (594 nm), a repetition rate of 40 MHz and a TCSPC time resolution of 16 ps. Fluorescence lifetime components were extracted from the low FRET efficiency (below 0.18) and the high FRET efficiency (above 0.75) populations. Photon weights were calculated for the two lifetime components (low FRET and high FRET population), according to Kapusta *et al*. (41). Subsequently, the same ClpB samples were measured at 1 nM, with pulsed excitation by the 485 nm laser at 50 µW, 40 MHz and TCSPC time resolution of 16 ps. Filtered FCS cross-correlation functions were calculated using the photon weights of the low FRET and high FRET populations as previously described (41). The filtered cross-correlation curves were fitted to the equation (Origin Pro 2019):

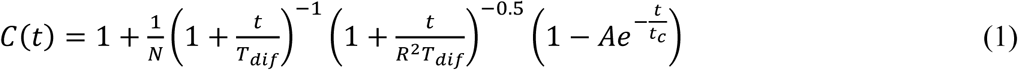

where is the number of molecules in the confocal volume, - diffusion time, - the ratio of the dimensions of the Gaussian-shaped beam waist parallel and perpendicular to the direction of light propagation, *A* is an amplitude and is the correlation time. The value of *R* was kept constant (at 6.2) for the fitting, and was determined from the FCS calibration measurement using a reference Rhodamine 6G solution with a manufacturer-provided diffusion coefficient (3.84×10^−6^ cm^2^s^-1^ in water at 22.4°C). The results are shown in Fig. 2c.

### Rotational simulation of the NTD

To obtain further information on the possible interactions of the NTD with the MD, we simulated multiple conformations of the NTD around its linker and mapped out possible interactions with the MD. First, we used the structural model of *TT* ClpB (6) to create multiple conformations of the NTD by rotating it as a rigid body around a point within the NTD-NBD1 linker (residue 142). We then excluded all conformations that caused a steric clash within the same protomer or with adjacent protomers. In our case, a clash was defined as a Van der Waals (VDW) contact between the MD atoms and the other atoms of the ClpB hexameric complex (equation 2):

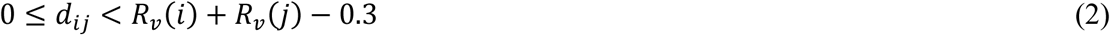

where *R*_*v*_(*i*) and *R*_*v*_ (*j*) are the VDW radii of atoms *i* and *j*, respectively, and *d*_*ij*_ is the distance between the two atoms. In equation 2, 0.3 Å is our clash threshold value (lower bound for a VDW contact), which was calculated from the structure of the hexameric model (37). We excluded all conformations that had at least one distance within the ranges calculated using equation 2. We then calculated the distances between all the NTD residues and the residue 487 on the MD. A threshold of 10 Å was used to select NTD structures within this range of distances. We found that α-helix A1 of the NTD was in close proximity to the MD (Fig. S3).

### Fluorescence measurements of Atto 655-labeled ClpB

Steady-state fluorescence of Atto 655 labeled ClpB mutants (either with no additional tryptophans (Ws), 12W or 23W) was measured using a Fluorolog-3 spectrofluorometer (Horiba Jobin Yvon, Edison, NJ, USA). Excitation wavelength was set to 635 nm, and fluorescence was collected between 650 and 800 nm. Measurements were conducted using 0.5 µM solutions of Atto 655-labeled ClpB mutants in 25 mM HEPES, 25 mM KCl, 10 mM MgCl_2_, 2 mM ATP, pH 8, at ambient temperature (25°C). All spectra were collected in duplicate per mutant, and background-corrected. Integrated average fluorescence values are listed in Table S4.

### Ensemble fluorescence lifetime measurements

Fluorescence lifetime measurements were performed on a FluoroHub time-correlated single-photon counting instrument (Jobin-Yvon), by using time-correlated single-photon counting method (46). 50 nM solutions of Atto 655-ClpB mutants, pre-filtered through 0.1 µm filters (Whatman Anotop-10), were measured (in 25 mM HEPES, 25 mM KCl, 10 mM MgCl_2_, 2 mM ATP, pH 8), with four repeats per sample. A pulsed diode laser with the emission at 635 nm was used as an excitation source, with a pulse length of 360 ps (FWHM). The emission was collected at 680 nm, with a slit-width of 12 nm. Photons were collected in 2048 channels, with a peak maximum of 10,000 counts. Data were globally fitted with 2-exponential model using a built-in fitting wizard, and average fluorescence lifetimes are reported (Table S5).

## Acknowledgements

M.I. is the recipient of an EMBO Long-Term Fellowship (ALTF 317–2018). H.M. was supported by Planning & Budgeting Committee of the Council of Higher Education of Israel. P.G. is funded by the Swiss National Science Foundation (Grant 31003A_156948). G.H. is funded by the European Research Council (ERC) under the European Union’s Horizon 2020 research and innovation programme (grant agreement No 742637), and is the incumbent of the Hilda Pomeraniec Memorial Professorial Chair. We are grateful to Dr. Yoav Barak for experimental support. We thank Dr. Rina Rosenzweig for ClpB plasmids and for excellent advice.

## Author contributions

M.I., H.M., I.R. and G.H. designed research; M.I., H.M. and P.G. performed research; M.I., H.M., P.G. analyzed data; M.I., H.M., I.R., P.G. and G.H. wrote the paper.

**The authors declare no conflict of interest**

## Supplementary Materials

Figs. S1-S9; Tables S1-S5; Supplementary Methods.

