## Supplementary Information for "Single-molecule spectroscopy reveals dynamic allostery mediated by the substrate-binding domain of a AAA+ machine"

### Supplementary Results

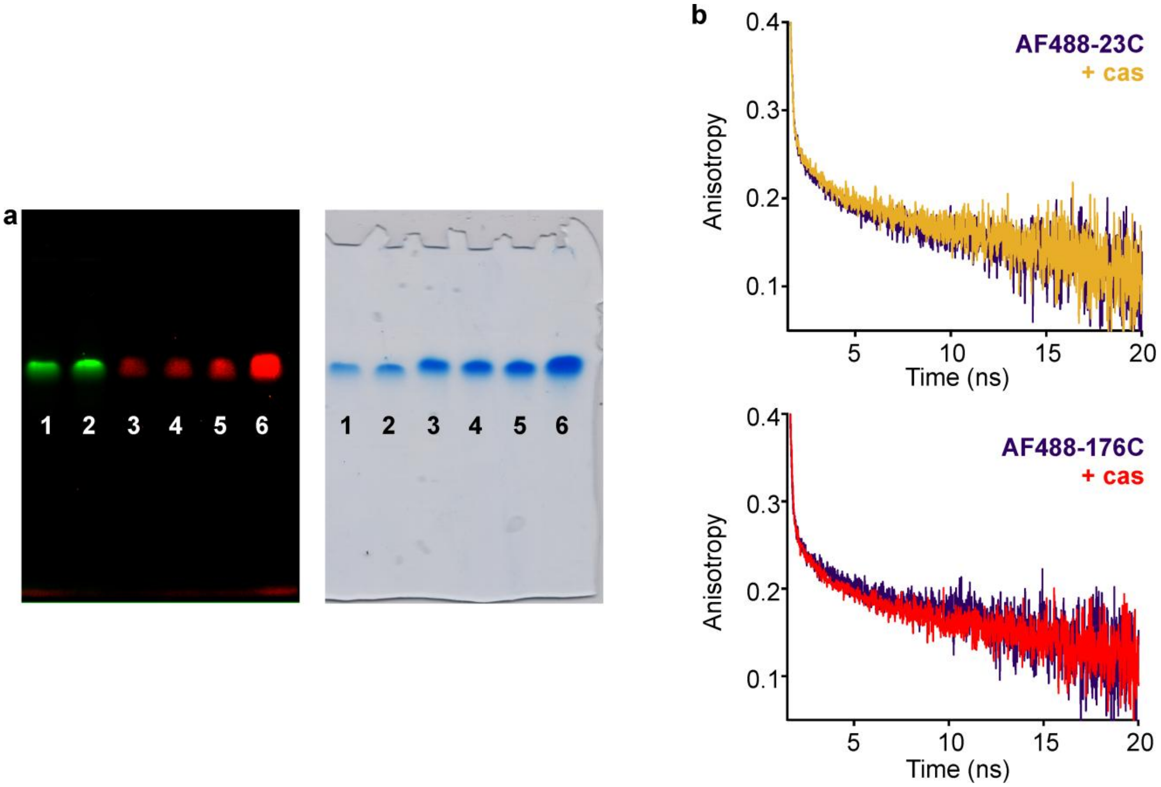


**Figure S1. Characterization of the 23C-176C mutant of ClpB, used to measure NTD dynamics. (a)** Representative native PAGE (6%) of fully fluorescently-labeled (with Alexa Fluor dyes (AF)) 23C-176C ClpB molecules in the presence of 2 mM ATP. Left- fluorescence image, right- white-light image after Coomassie staining. Lanes 1, 2- fully labeled AF488-23C-176C variant. Lanes 3-6- AF488 and AF594 double-labeled 23C-176C, mixed 1:100 with WT ClpB following a mixing by dialysis protocol detailed in the main text. Gel run for 12 h at 30 V and 4 ºC. **(b)** Time-resolved fluorescence anisotropy decay curves of labeled (with AF488) single-cysteine 23C and 176C mutants, mixed 1:100 with WT ClpB, and measured without and with 25 µM κ-casein ("+ cas"). The steady-state fluorescence anisotropy values, calculated as detailed in the Supplementary Methods section, were 0.205±0.001 (n=3) for AF488-23C, 0.201±0.001 (n=3) for AF488-23C with κ-casein, 0.208±0.001 (n=3) for AF488-176C and 0.206±0.001 (n=3) for AF488-176C with κ-casein. The errors correspond to standard errors of the mean. The initial fast decay of the anisotropy in all curves indicates free rotation of AF488 dyes attached to the residues 23C and 176C. The longer time anisotropy decay is due to the rotation of the protein as a whole. The similarity of the decays and the steady-state values with and without substrate protein indicates no influence of the latter on the freedom of motion of the dyes.

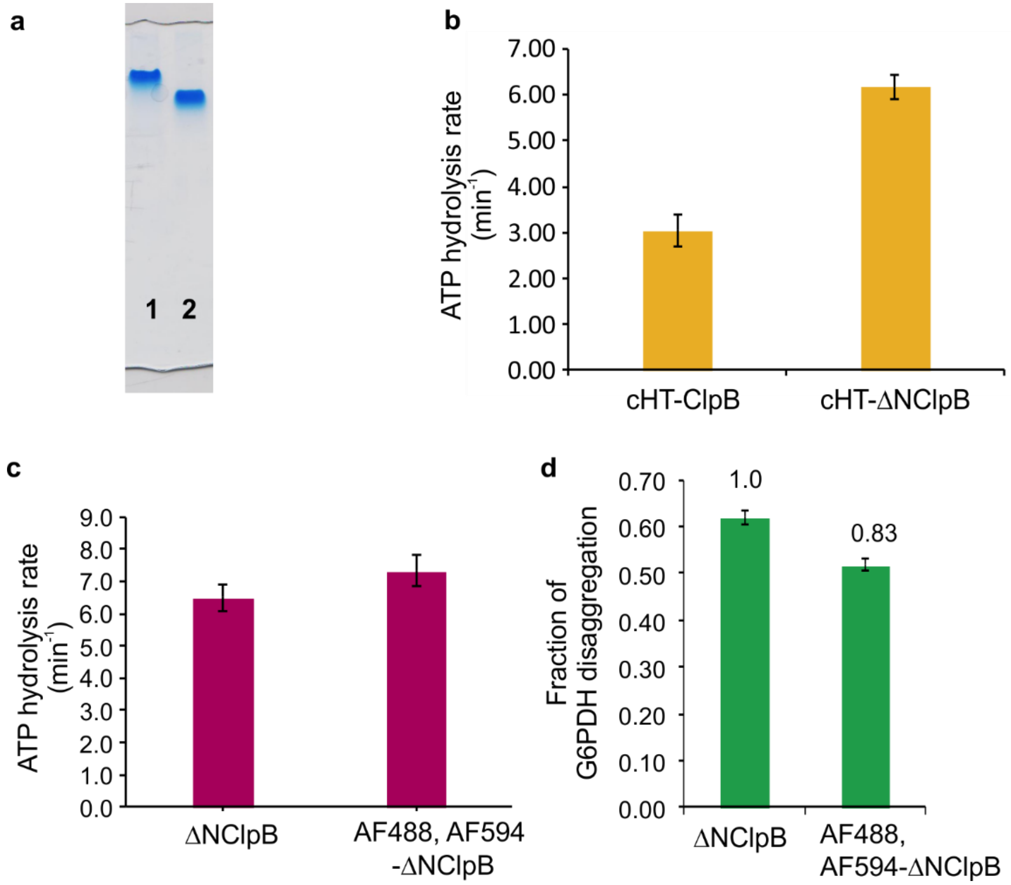


**Figure S2: Bulk experiments to test ATP activity and assembly of ClpB variants.** **(a)** Representative native gel (6%) of ClpB and ΔNClpB in the presence of 2 mM ATP. Lane 1- full-length ClpB, lane 2- ΔNClpB. **(b)** ATP activity measurement at 25°C in the presence of 2 mM ATP using ClpB and ΔNClpB proteins with cleaved six-histidine tag ("cHT"). The measured values were 3.1±0.5 (n=2) for ClpB and 6.2±0.4 (n=2) for ΔNClpB, in good agreement with the results for the proteins with an uncleaved six-histidine tag. (Fig. 3, main text). **(c)** Comparison of basal ATP activity at 25°C of unmodified unlabeled ΔNClpB and fully double-labeled mutant (AF488, AF594-) ΔNClpB (288C-631C). **(d)** Disaggregation of heat-induced aggregates of G6PDH, measured after 3 h of incubation in the presence of 2 µM ΔNClpB (either unmodified or fully double-labeled) with the addition of DnaK, DnaJ and GrpE (2 µM, 1 µM and 1 µM, respectively). The error bars correspond to standard errors of the mean.

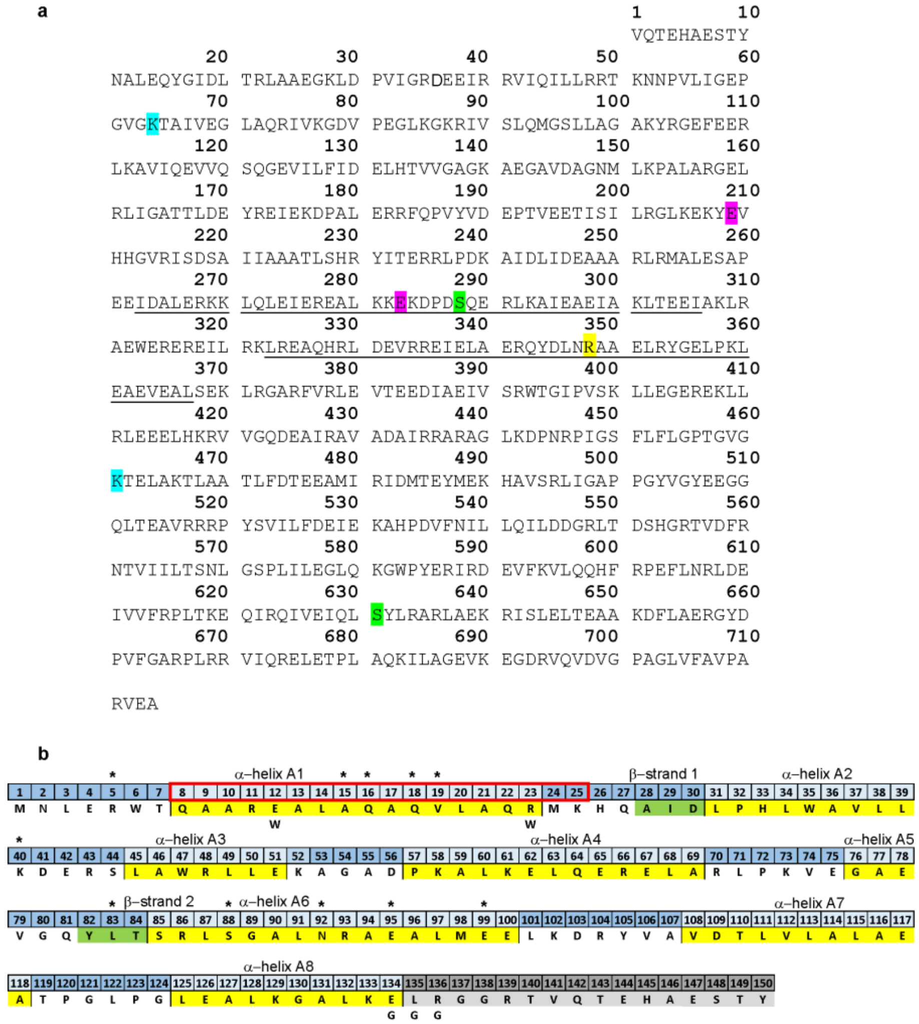


**Figure S3. Primary sequence of ΔNClpB and of the NTD. (a)** ΔNClpB construct used in this study (from *Thermus thermophilus)* begins with valine, corresponding to V141 in the full-length ClpB. The residues that were mutated are highlighted. Green: residues modified for AF dye incorporation, S288C (corresponding to S428C in full-length ClpB) and S631C (S771C in full-length ClpB). Purple: residues modified to alter middle domain (MD) dynamics. Activated mutation, E209A (E349A in full-length ClpB), and repressing mutation, E283A (E423A in full-length ClpB). Blue: residues modified to abolish ATP binding to either NBD1 or NBD2, Walker A1 mutation K64A (corresponding to K204A in full-length ClpB) and Walker A2 mutation K461A (K601A in full-length ClpB). Yellow: residue for Atto 655 incorporation R347C (R487C in full-length ClpB). **(b)** Primary sequence of the N-terminal domain (NTD) of full-length ClpB. Secondary structure elements are highlighted in yellow (α-helices) and in green (β-strands) (1). Flexible linker region connecting the domain to NBD1 is shown in grey. Residues found in proximity to the MD using geometrical simulation are marked with a black asterisk. Single amino-acid substitutions within the NTD, carried out in this study, are shown in bold letters below the sequence. Residues 8-25 (red box) were deleted in ΔHA1ClpB variant.

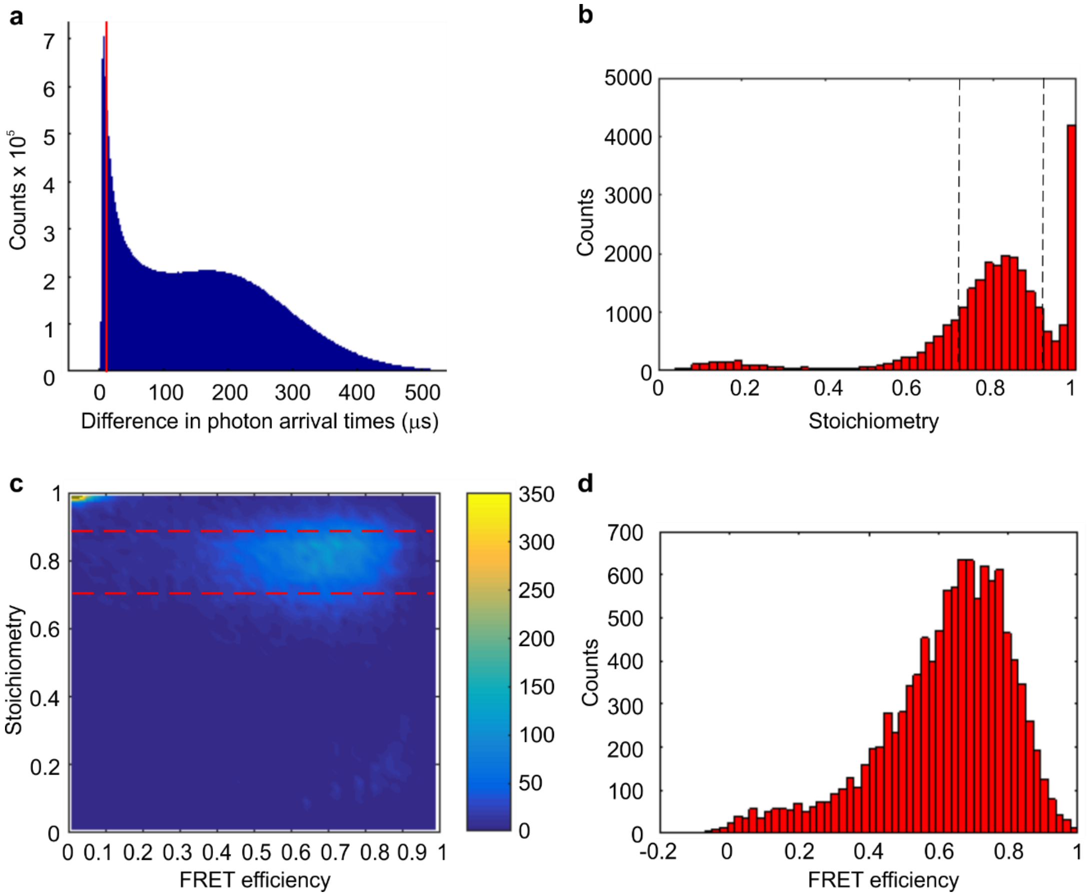


**Figure S4. Representative single-molecule results and data correction steps. (a)** Histogram of the time lags, recorded for the ΔNClpB (S288C-S631C) sample in the presence of 2 mM ATP. A cut-off time of 10 μs was used (red vertical line) to fully separate the photons emitted by the molecules from the background photons. **(b)** Representative stoichiometry histogram (50 bins), after correction for the leakage of photons from donor to acceptor channel. Stoichiometry, as described previously (2, 3), is computed as the ratio of the photons after donor (AF488) excitation to all photons after both donor and acceptor excitations, per fluorescence burst. Molecules bearing both the donor and acceptor dyes (AF488 and AF594, respectively) appear within the peak centered at the stoichiometry value of 0.83. Only molecules with the stoichiometry values between 0.73 and 0.91 are selected for further analysis to effectively eliminate any singly-labeled species. **(c)** Representative contour plot of stoichiometry versus FRET efficiency histogram (with 50 bins each). FRET efficiency is computed for each molecule after donor excitation only (as acceptor emission/(acceptor+donor emission)) (3). The dotted lines show the boundaries of selected stoichiometry values (0.73 and 0.91). FRET efficiency values of the molecules that have the stoichiometry values within this chosen range are selected. Then, only bursts with intensity greater than 30 photons are selected for further analysis. **(d)** The resulting FRET efficiency histogram (50 bins) for ΔNClpB (2 mM ATP) with 11,309 selected molecules. Subsequent $H^2$MM analysis, described in full detail previously (4), feeds in the information of individual photons emitted by the selected molecules (color and arrival times), and we use first 5,800 selected molecules from each dataset to ensure that the same number enters the analysis of the data recorded under various experimental conditions.

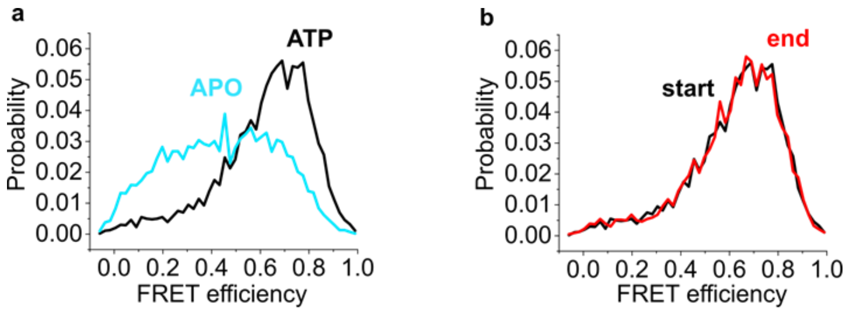


**Figure S5. FRET efficiency histograms from smFRET measurements of the MD dynamics of ΔNClpB. (a)** In the absence of ATP (blue), the FRET efficiency histogram is much broader than in its presence (black). This might be due to the partial disassembly of hexameric complexes in the absence of ATP, and therefore we carry out all smFRET experiments in the presence of 2 mM ATP. **(b)** FRET efficiency histograms derived from the measurement in the presence of 2 mM ATP. The acquired data are split in half, and FRET efficiency histograms for the first 100 min of detection (“start” in black) and the subsequent 100 min of detection (“end” in red) are compared. There is no change in their appearance, indicating that the analyzed species do not undergo changes during the measurement. All histograms contain 50 bins and are normalized to the total number of events.

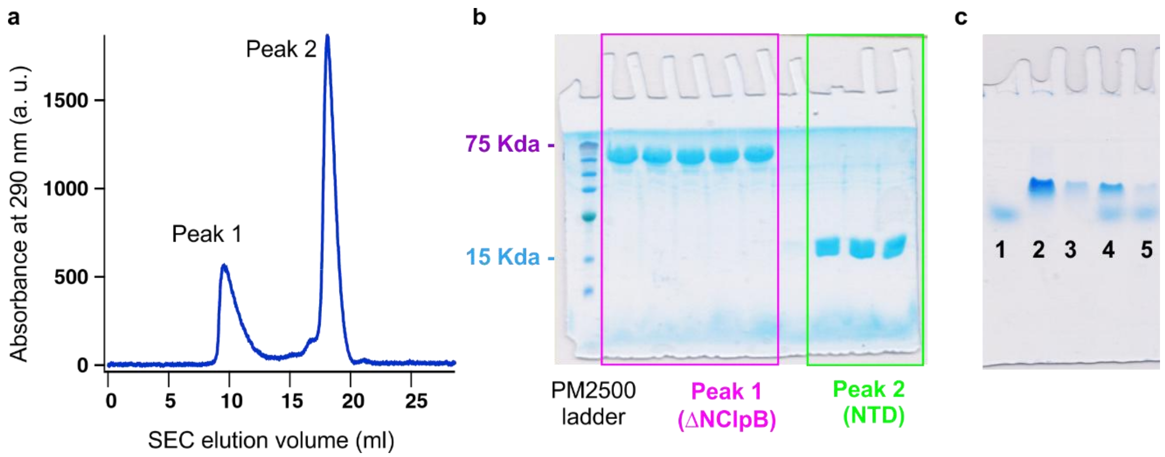


**Figure S6. Testing for the co-assembly of an isolated NTD of ClpB with ΔNClpB. (a)** Elution profile of a sample containing ΔNClpB (40 µM) and NTD (650 µM) (2 mM ATP, Superdex-200 column, GE Healthcare). Two high-intensity peaks are marked. **(b)** SDS-PAGE (15%) analysis of the two elution peaks. Peak 1 corresponds to ΔNClpB, and peak 2 is the NTD. Neither peak contains the second component, indicating that ΔNClpB and NTD elute separately. **(c)** Native PAGE of the NTD and ΔNClpB (6%, 2 mM ATP) shows that the two components migrate separately. The lanes are as follows: 1- NTD, 2- ΔNClpB; 3- ΔNClpB with cleaved six-histidine tag; 4- NTD+ΔNClpB; 5- NTD+ΔNClpB with cleaved six-histidine tag.

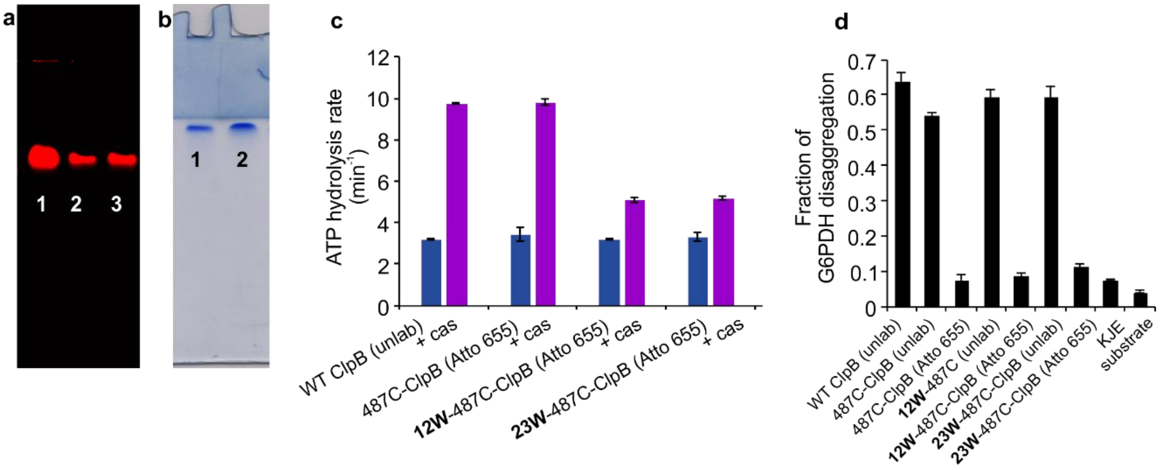


**Figure S7. Characterization of the mutants in the α-helix A1 of ClpB and of Atto 655-labeled mutants. (a)** Representative fluorescence image of the native PAGE (6%) of fully-labeled Atto 655-ClpB mutants in the presence of ATP (2 mM). Lane 1- Atto 655-ClpB; lane 2- 12W-Atto 655-ClpB, lane 3- 23W-Atto 655-ClpB. **(b)** Representative native gel (6%) confirming full assembly of ΔHA1 variant of ClpB (Δ8-25 aas) in the presence of 2 mM ATP. Lane 1- ΔHA1ClpB, lane 2- unmodified full-length ClpB. **(c)** ATPase activity of fully-labeled Atto 655-ClpB variants at 25 °C, without and with 50 µM κ-casein (n=2). **(d)** Disaggregation of the heat-induced aggregates of G6PDH as a substrate by 487C mutants of ClpB, either unlabeled (denoted as “unlab”) or fully-labeled with Atto 655 (denoted as “Atto 655”), measured after 3 h of incubation. ClpB mutants are at 2 µM, with the addition of DnaK, DnaJ and GrpE (2 µM, 1 µM and 1 µM, respectively). The fraction of disaggregation was determined by taking the amount of native (non-aggregated) G6PDH as 1 (n=4). The error bars correspond to standard errors of the mean.

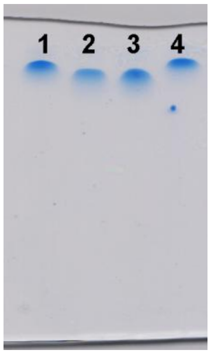


**Figure S8. Verification of the assembly of R136G variant.** Representative native PAGE (6%) in the presence of ATP (2 mM). R136G mutant migrates as a single band, similarly to the full-length ClpB. Both variants migrate above ΔNClpB. Lane assignment is as follows: 1- full-length ClpB, 2- ΔNClpB, 3- hyper-ΔNClpB; 4- full-length R136G mutant of ClpB.

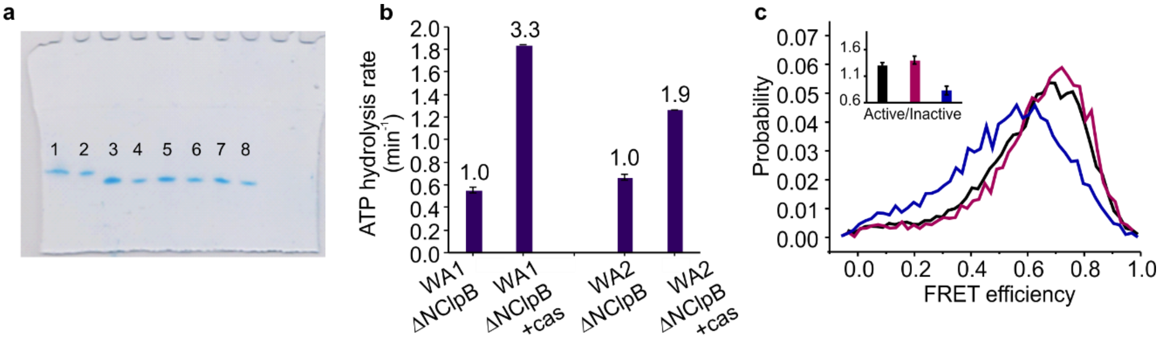


**Figure S9. Characterization of Walker A mutants of ΔNClpB. (a)** Representative native PAGE (6%) of unmodified unlabeled full-length ClpB, unmodified ΔNClpB, WA1-ΔNClpB (K64T, corresponding to K204T in the full-length ClpB) and WA2-ΔNClpB (K461T, corresponding to K601T in the full-length ClpB) variants in the presence of 2 mM ATP. Lanes 1, 2- ClpB (full-length), lanes 3, 4- ΔNClpB, lanes 5, 6- WA1-ΔNClpB, lanes 7, 8- WA2-ΔNClpB. All ClpB variants migrate as single bands, indicating correct assembly. **(b)** Rate of ATP hydrolysis of WA1-ΔNClpB and WA2-ΔNClpB mutants with and without the addition of 50 µM κ-casein at 25°C (n=2). **(c)** FRET efficiency histograms of Walker A mutants of ΔNClpB (S288C-S631C). Black- ΔNClpB, purple- WA1-ClpB, blue- WA2-ClpB. Active/inactive state ratios of the MD are shown in the inset. A prominent shift to lower FRET efficiency values for WA2-ΔNClpB indicates a higher proportion of the inactive state upon ATP binding to NBD1. Indeed, $H^2MM$ analysis confirmed this (Table S2, S3), showing that the MD in WA2-ΔNClpB mutant was repressed, with the corresponding ratio of active/inactive states of 0.83±0.08. Conversely, the MD in the WA1-ΔNClpB mutant was activated, with the active/inactive state ratio of 1.4±0.08, similarly to the value obtained for ΔNClpB. Thus, the suppression of the MD upon ATP binding to NBD1, seen in the full-length ClpB in our previous study (3), is unaffected by the NTD deletion. The error bars correspond to standard errors of the mean.

**Table S1: Analysis of ClpB's NTD dynamics. Comparison of the transition rates out of 3 states, derived from H²MM analysis and from dwell time distribution analysis** (3).

| Sample | H²MM analysis | | | Dwell time analysis | | |
|---|---|---|---|---|---|---|
| | **State** | **Rate (Hz)** | **SEM (n=4)** | **State** | **Rate (Hz)** | **SEM (n=3)** |
| 23C-176C | **1** | 3540 | 480 | **1** | 3650 | 590 |
| | **2** | 1860 | 150 | **2** | 1570 | 360 |
| | **3** | 2500 | 220 | **3** | 2450 | 70 |

| Sample | H²MM analysis | | | Dwell time analysis | | |
|---|---|---|---|---|---|---|
| | **State** | **Rate (Hz)** | **SEM (n=4)** | **State** | **Rate (Hz)** | **SEM (n=3)** |
| 23C-176C plus κ-casein (25 μM) | **1** | 3130 | 160 | **1** | 3190 | 120 |
| | **2** | 2570 | 60 | **2** | 2480 | 60 |
| | **3** | 3910 | 140 | **3** | 3970 | 210 |

**Table S2: Comparison of the relative occupancies of active (tilted) to inactive (horizontal) states of the MD**. Results of the H²MM analysis of ΔNClpB data, with 5,800 molecules per sample and 2-4 repeats per condition (SD is the standard deviation). The population ratio between active and inactive state is reported with the respective standard error of the mean (SEM).

| Sample | Inactive State | Inactive SD | Active State | Active SD | Active/Inactive | SEM |
|---|---|---|---|---|---|---|
| ΔNClpB | 0.44 | 0.02 | 0.56 | 0.02 | **1.30** | 0.05 |
| ΔNClpB plus κ-casein (25 μM) | 0.36 | 0.001 | 0.64 | 0.002 | **1.76** | 0.01 |
| Hyper-ΔNClpB | 0.35 | 0.01 | 0.65 | 0.01 | **1.85** | 0.07 |
| Repr-ΔNClpB | 0.57 | 0.04 | 0.43 | 0.04 | **0.76** | 0.07 |
| WA1-ΔNClpB | 0.42 | 0.02 | 0.58 | 0.02 | **1.40** | 0.08 |
| WA2-ΔNClpB | 0.45 | 0.02 | 0.55 | 0.02 | **0.83** | 0.08 |
| ΔHA1ClpB | 0.43 | 0.01 | 0.57 | 0.01 | **1.32** | 0.06 |
| Full-length R136G | 0.47 | 0.01 | 0.53 | 0.01 | **1.14** | 0.02 |
| Full-length ClpB (3) | 0.49 | 0.01 | 0.51 | 0.01 | **1.00** | 0.01 |

**Table S3: Comparison of the transition rates between inactive and active states of the MD, derived from $H^2MM$ analysis.**

| Sample | Active to Inactive, $k_{12}$ (Hz) | SEM | Inactive to Active, $k_{21}$ (Hz) | SEM |
|---|---|---|---|---|
| ΔNClpB | 4780 | 217 | 6296 | 92 |
| ΔNClpB plus κ-casein (25 μM) | 5278 | 322 | 9573 | 517 |
| Hyper-ΔNClpB | 4123 | 375 | 7840 | 382 |
| Repr-ΔNClpB | 5476 | 146 | 4154 | 269 |
| WA1-ΔNClpB | 4551 | 193 | 6527 | 134 |
| WA2-ΔNClpB | 3927 | 249 | 3259 | 423 |
| ΔHA1ClpB | 4397 | 111 | 5917 | 310 |
| Full-length R136G | 4446 | 88 | 5255 | 110 |
| Full-length ClpB (3) | 5300 | 150 | 5700 | 100 |

**Table S4: Results of the steady-state fluorescence measurements of Atto 655-labeled ClpB mutants.**

| Sample | Fluorescence Intensity (a. u.) | SD (n=2) |
|---|---|---|
| Atto 655-487C ClpB | $8.08 \times 10^7$ | $2.92 \times 10^5$ |
| 12W Atto 655-487C ClpB | $2.93 \times 10^7$ | $2.12 \times 10^2$ |
| 23W Atto 655-487C ClpB | $3.15 \times 10^7$ | $8.06 \times 10^3$ |

**Table S5: Results of the fluorescence lifetime measurements of Atto 655-labeled ClpB mutants.**

| Sample | Lifetime (ns) | SD (n=4) |
|---|---|---|
| Atto 655-487C ClpB | 2.04 | $4.07 \times 10^{-3}$ |
| 12W Atto 655-487C ClpB | 1.29 | $1.93 \times 10^{-3}$ |
| 23W Atto 655-487C ClpB | 1.32 | $9.00 \times 10^{-3}$ |

### Supplementary Methods

**Protein expression and purification** was performed closely following the recently reported protocols (3). *Thermus thermophilus* ClpB (*TT* ClpB) or ΔNClpB (*TT* ΔNClpB) DNA was cloned into a pET28b vector with kanamycin resistance, and a six-histidine tag was added at the start of the sequence, preceded by a tobacco etch virus protease (TEV) cleavage site. All mutations in ΔNClpB were introduced using standard site-directed mutagenesis and confirmed by DNA sequencing. *E. coli* BL21 (DE3) cells were transformed with the ClpB vector and grown at 37°C with shaking to OD 0.7-0.8 in the presence of kanamycin (Caisson Laboratories). Protein expression was induced by adding 1 mM IPTG, followed by an overnight incubation with shaking at 25 °C. Subsequently, bacteria were harvested and the protein was purified on a Ni-NTA resin (GE Healthcare) and eluted with 250 mM imidazole (Sigma Aldrich). This was followed by an overnight dialysis at 4°C in the presence of 2 mM ATP (25 mM HEPES, 25 mM KCl, 2 mM TCEP, 10 mM $MgCl_{2,}$ 2 mM ATP) to remove imidazole from the solution. Following filtration (0.22 µm Millex, Merck), the protein was further purified using a HiPrep DEAE FF column (GE Healthcare) equilibrated with 50 mM HEPES, 20 mM KCl and 2 mM TCEP at pH 7.4. The peak containing the purified protein was collected, aliquoted, flash-frozen and stored at -80 °C. The purity was verified by gel electrophoresis. The aliquots were thawed immediately prior to experiments and used once. Expression and purification of any ClpB mutational variants, DnaK, DnaJ and GrpE followed similar procedures. Histidine-tag removal was carried out by adding ~6 mg histidine-tagged TEV protease to 20 mg protein during an overnight dialysis step after the first Ni-NTA column. The cleaved protein was then separated from the protease and any uncleaved protein on a Ni-NTA column, and the subsequent purification steps were performed as described.

**Fluorescence anisotropy decay measurements** were carried out using single-cysteine mutants of ClpB, 23C and 176C, labeled with AF488 and mixed with 100-fold molar excess of unmodified WT ClpB using the mixing by dialysis procedure described in the Methods section (main text). The samples were diluted to 1 nM (in 25 mM HEPES, 25 mM KCl, 10 mM $MgCl_2$, 2 mM ATP, 0.01% TWEEN 20), with and without 25 µM κ-casein, filtered (0.1 µm filters, Whatman Anotop-10) and loaded into the flow chambers prepared according to the same protocol as for single-molecule measurements (main text). Data were collected using a MicroTime200 fluorescence microscope (PicoQuant), with 485 nm laser excitation (10 µW), 20 MHz repetition rate and 16 ps time resolution. Fluorescence was passed through a 50 µm pinhole, split into parallel and perpendicular components using a polarizing beam splitter (Ealing), filtered through band-pass filters (520/35 nm, Semrock) and collected by two single-photon avalanche photodiode detectors (Excelitas SPCM-AQR-14-TR) coupled to a time-correlated single-photon counting module (HydraHarp 400, PicoQuant). Time-resolved fluorescence anisotropy decays were then calculated as follows:

$$r(t) = \frac{I_{\parallel(t)} - I_{\perp(t)}}{I_{\parallel(t)} + 2I_{\perp(t)}}$$

where $I_{\parallel(t)}$ and $I_{\perp(t)}$ are the time-dependent fluorescence intensities of parallel and perpendicular components, respectively. Steady state fluorescence anisotropy was calculated using the integrated signal according to:

$$r = \frac{\sum I_{\parallel(t)} - \mathrm{G} \sum I_{\perp(t)}}{\sum I_{\parallel(t)} + 2\mathrm{G} \sum I_{\perp(t)}}$$

where G is the polarization sensitivity factor (whose value was 1 in our setup). The results are shown in Fig. S1.

**Disaggregation of heat-induced aggregates of G6PDH.** For the preparation of glucose-6-phosphate dehydrogenase (G6PDH) aggregates, 90 µM G6PDH from *Leuconostoc mesenteroides* (Sigma Aldrich) was denatured for 5 min at 47°C in the presence of 5 M urea, 20 mM DTT and 7.5% glycerol. Then it was diluted 1:100 into a reactivation buffer (50 mM HEPES, 20 mM $MgCl_2$, 30 mM KCl, 1 mM EDTA, 1 mM TCEP, 3 mM ATP and 20 µg/µl pyruvate kinase) and further incubated at 47°C for 15 min. For the disaggregation assay, 750 nM of the aggregated G6PDH was combined with 2 µM ClpB (or with 2 µM ClpB variants), 2 µM DnaK, 1 µM DnaJ and 1 µM GrpE in the reactivation buffer, and incubated at 37°C for up to 3 hours. The activity of soluble G6PDH was measured both before and after the incubation with ClpB mixture, by withdrawing aliquots and diluting them into activity buffer (5 mM $MgCl_2$, 2 mM D-glucose-6-phosphate and 1 mM $NADP^+$). The activity was measured by monitoring the time-dependent linear increase in absorption at 340 nm due to NADPH formation. The steepest linear increase in absorption was observed for native non-heated G6PDH. The fraction of disaggregation was reported as the slope measured for the samples divided by the slope measured for native G6PDH. To obtain the concentration of the recovered G6PDH, the fraction of disaggregation was multiplied by the starting concentration (750 nM). To obtain the disaggregation rates, we measured the activity of soluble G6PDH in the disaggregation mixtures over time, and plotted the concentration of recovered G6PDH as a function of time during the incubation at 37°C. The disaggregation rate was calculated as a slope of the initial linear part of the curve (first 50 min).
